# Conserved whole-brain spatiomolecular gradients shape adult brain functional organization

**DOI:** 10.1101/2022.09.18.508425

**Authors:** Jacob W Vogel, Aaron Alexander-Bloch, Konrad Wagstyl, Maxwell Bertolero, Ross Markello, Adam Pines, Valerie J Sydnor, Alex Diaz-Papkovich, Justine Hansen, Alan C Evans, Boris Bernhardt, Bratislav Misic, Theodore Satterthwaite, Jakob Seidlitz

## Abstract

Cortical arealization arises during neurodevelopment from the confluence of molecular gradients representing patterned expression of morphogens and transcription factors. However, how these gradients relate to adult brain function, and whether they are maintained in the adult brain, remains unknown. Here we uncover three axes of topographic variation in gene expression in the adult human brain that specifically capture previously identified rostral-caudal, dorsal-ventral and medial-lateral axes of early developmental patterning. The interaction of these spatiomolecular gradients i) accurately predicts the location of unseen brain tissue samples, ii) delineates known functional territories, and iii) explains the topographical variation of diverse cortical features. The spatiomolecular gradients are distinct from canonical cortical functional hierarchies differentiating primary sensory cortex from association cortex, but radiate in parallel with the axes traversed by local field potentials along the cortex. We replicate all three molecular gradients in three independent human datasets as well as two non-human primate datasets, and find that each gradient shows a distinct developmental trajectory across the lifespan. The gradients are composed of several well known morphogens (e.g., *PAX6* and *SIX3*), and a small set of genes shared across gradients are strongly enriched for multiple diseases. Together, these results provide insight into the developmental sculpting of functionally distinct brain regions, governed by three robust transcriptomic axes embedded within brain parenchyma.

## Introduction

The brain coordinates different facets of behavior through diverse cell populations that are spatially distinct yet anatomically connected through a network of white matter fibers. This topographical distribution of function in the brain emerges during early development through the process of arealization, where cell differentiation and migration create a mosaic of brain areas with distinct molecular properties (Cadwell et al., 2019; Hébert and Fishell, 2008). One of the key tenets of arealization is the expression of morphogen gradients along the cardinal axes of the developing brain. Morphogens are produced from vesicular patterning centers in the notochord and diffuse along developing neural compartments, forming semi-orthogonal concentration gradients as the forebrain emerges (Bishop et al., 2000; O’Leary et al., 2007; Rakic, 1988). Morphogen expression triggers a cascade of molecular differentiation leading to the formation of spatially distributed gradients (Ypsilanti et al., 2021). These large-scale molecular gradients establish an organizational scaffold where areas distant from one another are also distinct in their molecular (and emergent functional) properties (Cadwell et al., 2019).

Molecular gradients have been previously described in the developing brain, namely those that radiate along rostral-caudal (longitudinal), dorsal-ventral (vertical) or medial-lateral (horizontal) axes of the neural compartment (Grove and Fukuchi-Shimogori, 2003; Hoch et al., 2009). Confluent spatial expression of these gradients in turn define important divisions within the brain along those same axes, which appear to be strongly conserved across phylogeny (Meinhardt, 2008). Cerebral organoids, which model early prenatal development (Lancaster et al., 2013), will spontaneously self-organize along similar spatial axes (Kadoshima et al., 2013). Furthermore, the establishment of molecular patterning gradients appears to be a critical component of normal brain development, as variations to genes regulating these gradients can cause developmental malformations, many of which cause severe morbidity or mortality (Flores-Sarnat and Sarnat, 2007).

Recent work has also focused on how molecular gradients may help to define the functional organization of the cerebral cortex, which exhibits substantially less molecular differentiation and a more uniform physiology compared to other parts of the brain (Hawrylycz et al., 2012). Nonetheless, the cerebral cortex exhibits functionally distinct territories with subtle variations in physiology (Glasser et al., 2016), and can be further subdivided into smaller regions with ostensibly differentiated roles in cognition and behavior (Eickhoff et al., 2018). A robust functional hierarchy has been described that situates different cortical brain regions along a sensorimotor-association axis (Bernhardt et al., 2022; Burt et al., 2018; Harris et al., 2019; Margulies et al., 2016; Sydnor et al., 2021). Besides discriminating functional roles of different cortical territories, variation in other physiological properties of the cortex occur along a similar topography to the functional hierarchy (Burt et al., 2018; Fulcher et al., 2019; Gao et al., 2020; Wang, 2020), and it may serve as a principal avenue for directional functional signals (Manea et al., 2022; Pines et al., 2022; Siegle et al., 2021; Vézquez-Rodríguez et al., 2020). Cortical peaks in the functional hierarchy are distributed nearly maximally distant from one another (Margulies et al., 2016; Oligschläger et al., 2019), suggesting that the functional hierarchy may arise due to interacting underlying transcriptomic gradients (Wagstyl et al., 2022). Furthermore, several studies have used transcriptomic data densely sampled across the adult cortex to show principal components of cortical gene expression, and have noted their topographic similarity to the functional hierarchy (Anderson et al., 2020; Burt et al., 2018; Hawrylycz et al., 2015). What is not entirely clear is whether and how the morphogenetic gradients that give rise to early developmental patterning relate to the adult functional hierarchy.

Although previous studies have identified heterochronicity in the spatiotemporal expression of many genes in the human brain (Li et al., 2018; Somel et al., 2009), prior work has not investigated whether developmental directional gradients of gene expression are still present in adulthood, and whether or how they relate to functional specialization. Regional development of the cerebral cortex appears to occur along specific directional gradients in terms of peak growth and rate of change (Bethlehem et al., 2022). Furthermore, heritability analyses suggest a genetic basis to the organization of cortical surface area and thickness along principal anterior-posterior and dorsal-ventral aspects, respectively (Chen et al., 2012, 2013; Valk et al., 2020). Follow up work has suggested that these genetically imposed patterns are disrupted in developmental disorders (Ball et al., 2020; Lombardo et al., 2021). More recent complementary work has provided evidence for an isolated directional gradient of gene expression in the adult brain, relating this gradient to other brain (neurodegenerative) diseases (Vogel et al., 2020). Together, these studies suggest directional developmental molecular gradients may be present in the adult brain, are distinct from the functional hierarchy, and may be relevant for brain-related traits and disorders. However, whether and how transcriptional gradients in the adult brain relate to molecular gradients underlying early fetal brain development remains unclear.

In the present study, we used a multivariate data-driven approach to establish directional gradients of gene expression in the adult human brain that vary along the axes of morphogen gradients in the developing brain. Unlike previous studies, we sample directional gene expression across the entire brain, not just the cerebral cortex, to match the propagation of gradients through the developing fetal brain. We validate the presence of identified gradients in three independent datasets of postmortem adult human tissue, track their evolution over the course of development, and examine their phylogenetic stability by examining the presence of these gradients in the brains of non-human primates. Importantly, we show that whole-brain gradients converge in the cerebral cortex to form recognizable divisions representing established cytoarchitectonically-defined cortical territories. Finally, we show that genes involved in the coordination of multiple gradients are specifically associated with brain diseases, underscoring the importance of molecular gradients as organizational foundations for the healthy brain. Together, our results delineate conserved developmental gradients that are transcriptomically encoded in the adult brain, and which represent an important component of adult cortical differentiation.

## Results

### Spatial transcriptomic gradients in the adult human brain

We hypothesized that large-scale developmental transcriptomic gradients (**Fig 1A**) would be detectable in the adult human brain as directional autocorrelation of gene expression spanning the entire brain. To delineate adult transcriptomic gradients, we cross-decomposed a correlated expression matrix composed of 15,634 genes across 3,466 tissue samples (see Methods) with a matrix of three-dimensional Euclidean coordinates for the same tissue samples (**Fig S1A**). The resulting components represent the principal sources of directional linear variation in whole-brain gene expression. A three-component partial least squares (PLS) regression (Wold S, 2001) model provided the best reconstruction of sample location based on repeated 10-fold cross-validation on training data (70% of samples) (**Fig S1C; 1B**). Tissue expression across the three transcriptomic PLS latent variables (LV1, LV2, LV3) explained the majority of variance across all three Euclidean coordinates of left out tissue samples (X R^2^=0.55, Y R^2^=0.69, Z R^2^=0.82) and was sufficient to predict the location of left out tissue samples (test data; 30% of samples) extracted throughout the brain with an average of 11.4 mm error (**Fig 1E, Fig 1H**). The composition of the gene expression LVs were also largely invariant to spatial permutations of the brain (**Fig S1F**).

Having demonstrated robust generalizability, the PLS model was then fitted using all data, and was used to reconstruct the location of all tissue samples (**Fig 1F,G**; see **Table S1** for component-weighted genes). Based on these findings, the PLS latent variables can collectively be considered as a set of three-dimensional spatiomolecular axes along which brain regions are putatively organized or embedded. Notably, reconstruction was least accurate for tissue samples extracted from the polar extremes of frontal, parietal and temporal cortex (**Fig 1H; Fig S2**). While this phenomenon may be a byproduct of linear modeling, it may also relate to the relatively limited variance of gene expression across cortical areas (Hawrylycz et al., 2012; Li et al., 2018), or to the extreme expansion of these regions among primates (Hofman, 2014); these three scenarios are not mutually exclusive.

**Figure 1:**
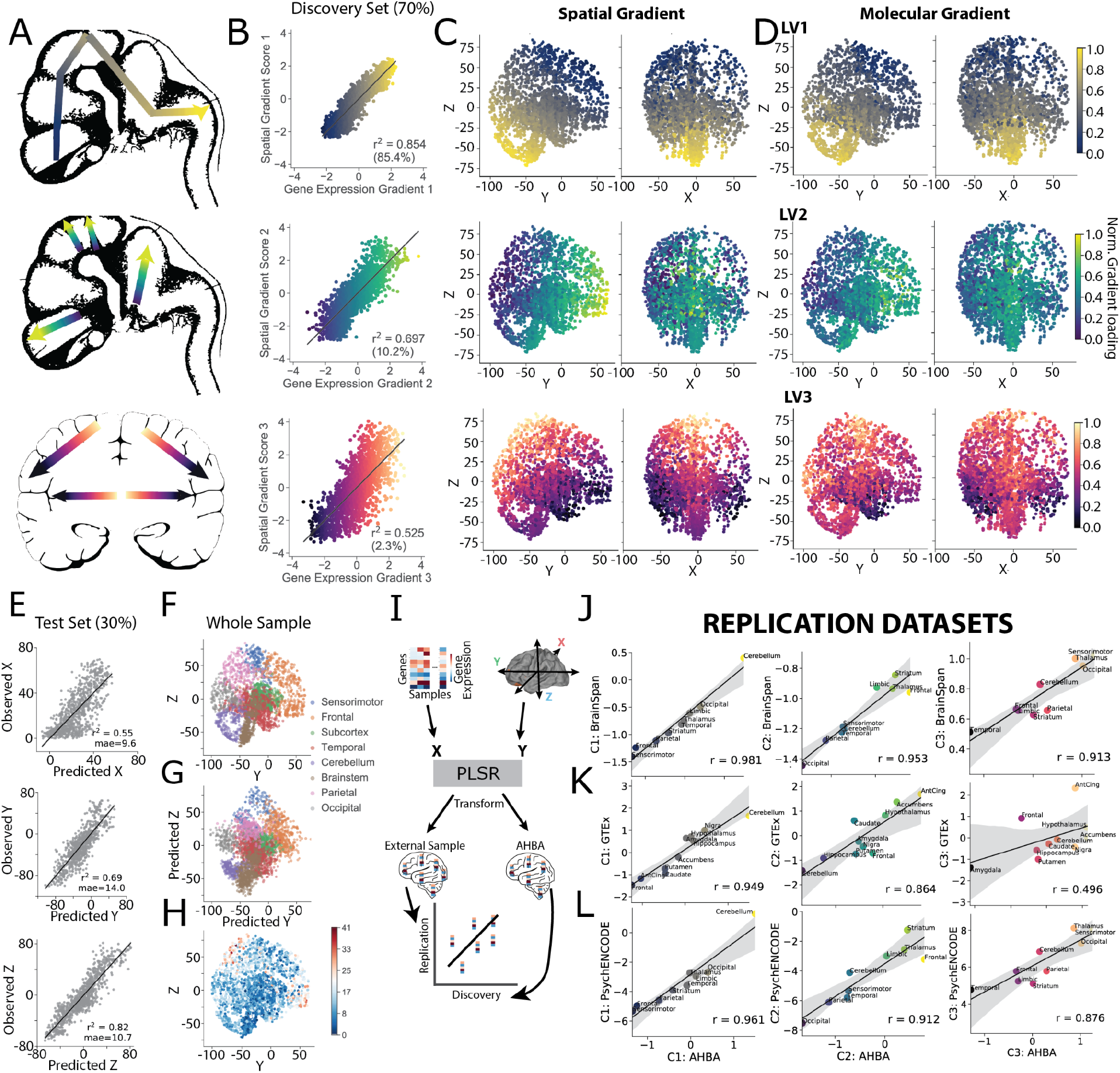
Large-scale spatiomolecular axes of the adult human brain. Partial Least Squares (PLS) analysis finds strong covariance between gene expression and three-dimensional spatial coordinates, summarized by three components resembling previously described whole-brain developmental molecular gradients. **A)** Theoretical whole-brain molecular gradients of the developing human brain, adapted from Flores-Sarnat & Sarnat (2008). This includes (top) a rostral-caudal gradient extending between forebrain and hindbrain, (middle) a dorsal-ventral gradient traversing each compartment, and (bottom) dorsomedial-ventrolateral (horizontal) gradient in the telencephalon. **B)** A PLS regression model was used to link whole-brain gene expression with Euclidean spatial coordinates in a training sample representing 70% of the total dataset (see Methods, **Figure S1A**). This model derived gradients of systematic linear variation in gene expression across spatial axes. Scatter plots show each of the three PLS LVs in the form of correlations between latent spatial and transcriptomic variables. **C)** Representation of the latent spatial variables of, from left-to-right, LV1, LV2 and LV3. These latent variables represent the main linear axes along which gene expression covaries through space. **D)** Representation of the latent transcriptomic variables of LV1, LV2 and LV3. These latent variates show the matching patterns of gene expression that vary along axes defined in C). Note that correlations between C) and D) are represented by the scatter plots in B). **E)** The three latent PLS variables were together able to predict the spatial coordinates (from top-bottom, x, y and z) of unseen tissue samples based on each sample’s gene expression data alone.The unseen samples consisted of 30% of tissue samples left out of the PLS model fitting. **F)** Y and Z coordinates of all Allen Brain Atlas tissue samples. **G)** PLS-predicted Y and Z coordinates of the same samples (see Supp File for 3D view). **H)** Plot showing distribution of prediction error in space (average of 11.4mm). **I)** The PLS model was refitted using only genes shared across discovery and replication datasets to allow us to test the reliability of the three spatiomolecular axes in entirely new data. This model was used to transform gene expression from the discovery and replication tissue samples into latent transcriptomic variables that compose the PLS LVs. Regional expression of each component in the discovery dataset was correlated against regional expression in the replication dataset. The observed regional correlation between AHBA PLS LVs and external PLS LVs indicates the degree to which regional expression of each component is similar across datasets. A high correlation indicates a similar gradient across a similar spatio-molecular axis exists in the external data. Dataset correlation is averaged over adults from the **J)** Brainspan (n=6), **K)** GTEx (n=121), and **L)** PsychENCODE (n=6) datasets. The observed spatio-molecular axes were highly consistent across datasets with the exception of LV3 expression in GTEx.

Despite being generated using a data-driven approach, the three PLS latent variables closely resembled theorized developmental whole-brain spatial gene expression gradients (**Fig 1A,C,D**). These patterns could not be explained by regional variation in aggregate gene expression across samples (**Fig S3B**).The first component (LV1) featured gene expression varying along a rostro-caudal axis, radiating from the brainstem and cerebellum, through the midbrain, subcortex and posterior cortex, and finally into anteriodorsal cerebral cortex. Expression of this component demonstrated linear rostro-caudal increase as one moves along regions originating from different segments of the neural tube (**Fig S3C**). The second component (LV2) was expressed in a whole-brain dorsal-ventral pattern, with highest expression in the brainstem, subcortex and frontal cortex, and lowest expression in cerebellar and posterior cerebral cortex. Within regions originating from neural tube segments, LV2 expression differed between regions originating from the dorsal and ventral plates (**Fig S3D**). The third component (LV3) described gene expression varying along a gradient diffusing in a medial-lateral and dorsal-ventral direction. Of the three gradients, the LV3 gradient showed the most variation within the cerebral cortex (**Fig S3A**). The cortical expression of the LV3 gradient also bore a strong relationship to the pattern of cortical myelination, which has been shown to be a robust index of hierarchical functional organization (Burt et al., 2018; Foit et al., 2022) (r=0.78, p<0.001; **Fig S3E**).

We hypothesized that cortical organization may be influenced by whole-brain spatial expression gradients above and beyond intracortical gradients. Interestingly, each PLS latent variable had a unique distribution of cortical expression (**Fig S3F**). Low LV1 expression differentiated the cerebral cortex from the rest of the brain. Meanwhile, LV2 expression was high in non-cortical structures, strongly differentiating it from cortex (notably including cerebellar cortex). Finally, LV3 expression was highest particularly in different types of sensory cortex, differentiating it from other parts of the brain, including association cortex. These analyses demonstrate how association cortex, sensory cortex, cerebellar cortex, and non-cortical structures show distinct patterns of spatiomolecular organization in the human brain.

### Transcriptomic gradient expression is consistent across adult human datasets

While spatially comprehensive, the discovery dataset for the three transcriptomic gradients was composed of gene expression data from only six individuals. Therefore, we sought to test the degree to which expression of these gradients generalized to other datasets of brain gene expression. We used the PLS model trained in the discovery dataset (AHBA) to transform the expression of the molecular gradients in the other replication datasets, BrainSpan and PsychENCODE (**Fig 1I**). We verified the efficacy of these predictions by validating that the gradients had similar expression profiles across spatially distributed brain regions. We found excellent reproducibility of all three gradients when averaging across adult individuals, despite different brain regions being measured across datasets (**Fig 1J,K,L**). In general, replication of both LV1 (range: r=0.95-0.98) and LV2 (r=0.86-0.95) were quite strong, while LV3 replication was more variable (r=0.50-0.91). In the Brainspan and PsychENCODE datasets, comparisons had consistent effect sizes when averaging across individuals of all ages rather than just adults (**Table S2**). In the GTEx sample, the effect sizes remained stable whether or not including individuals over 60 or including individuals with evidence for brain disease or other issues that might confound transcriptomic data (**Table S2**).

### Whole-brain spatial gradients capture the patterning of biological features across the cerebral cortex

The above findings led us to speculate that whole-brain spatial gradients may be associated with other fundamental properties of cortical organization. One common way to understand cortical functional organization is through primary gradients of functional connectivity variance derived from fMRI, which are thought to define multiple cortical functional hierarchies (Margulies et al., 2016). We directly compared the explanatory power of functional connectivity gradients to that of our spatial transcriptomic gradients by building models composed of each (and both) of these gradients to predict several brain features (**Fig 2A,B,C**). We found that molecular cortical gradients (**Fig 2A**) explained significantly more variance than functional gradients in eight of 11 brain features assessed, including cortical thickness and myelination, aerobic glycolysis, and allometric scaling (**Fig 2B**). In contrast, functional gradients explained more variance than the molecular gradients only in meta-analytic functional coactivation patterns. We next conducted models using both types of gradients to discern whether the molecular and functional gradients were contributing overlapping or distinct information to the spatial topography of cortical features (**Fig 2C)**. Models for five variables (aerobic glycolysis, geometric distance, principal meta-analytic functional gradient, intrinsic timescale and principal gene-related functional gradient) explained significantly more variance when both types of gradients were included, indicating that functional and molecular modalities contributed complementary information. This was most striking for aerobic glycolysis, for which 75% of the variance in spatial topography could be explained by the combined gradients. These analyses together delininate a strong relationship between variation in fundamental properties of brain organization and the distribution of spatiomolecular gradients.

Given the causal role of interacting gene expression gradients in the developmental process of arealization, we hypothesized that cortical overlap of whole-brain spatiomolecular gradients may be involved in macroscale organization of regions with distinct areal boundaries. To evaluate this, we performed a clustering analysis to identify distinct territories of spatiomolecular gradient that overlap in the cerebral cortex, which converged on six distinct territories (**Fig 2E, S4A,B**). The spatial extent of the territories overlapped with established functional and cytoarchitectonic subdivisions of the cerebral cortex: Territory 1 (blue) outlined motor and premotor cortex, Territory 2 (green) covered the prefrontal cortex, Territory 3 (purple) included parietal and lateral occipital cortical areas, Territory 4 (pink) subsumed limbic and subcortical regions, Territory 5 (yellow) outlined primary visual cortex, and Territory 6 (teal) encompassed the temporal lobe (**Fig 2E, Fig S4B**). The molecular territories were compared directly to a lobar parcellation of the cortex, and the two parcellations showed good agreement (adjusted mutual information score = 0.57, p_spin_<0.001, using 1,000 spatially-preserving permutations; adjusted Rand index = 0.46, p_spin_<0.001; **Fig 2E**). To further explore whether these territories indeed represent functional divisions of the telencephalon, we used meta-analytic functional coactivation data (Yarkoni et al., 2011) to examine the behaviors associated with activation of regions within each territory (**Fig S4C**). This analysis recapitulated well established structure-function relationships in the brain. Namely, the six cortical territories were differentially involved in tasks and movement (Territory 1; blue), social cognition (2; green), spatial orientation (3; purple), valuation and encoding (4; pink), visual function (5; yellow), and language (6; teal) (**Fig S4C**).

**Figure 2:**
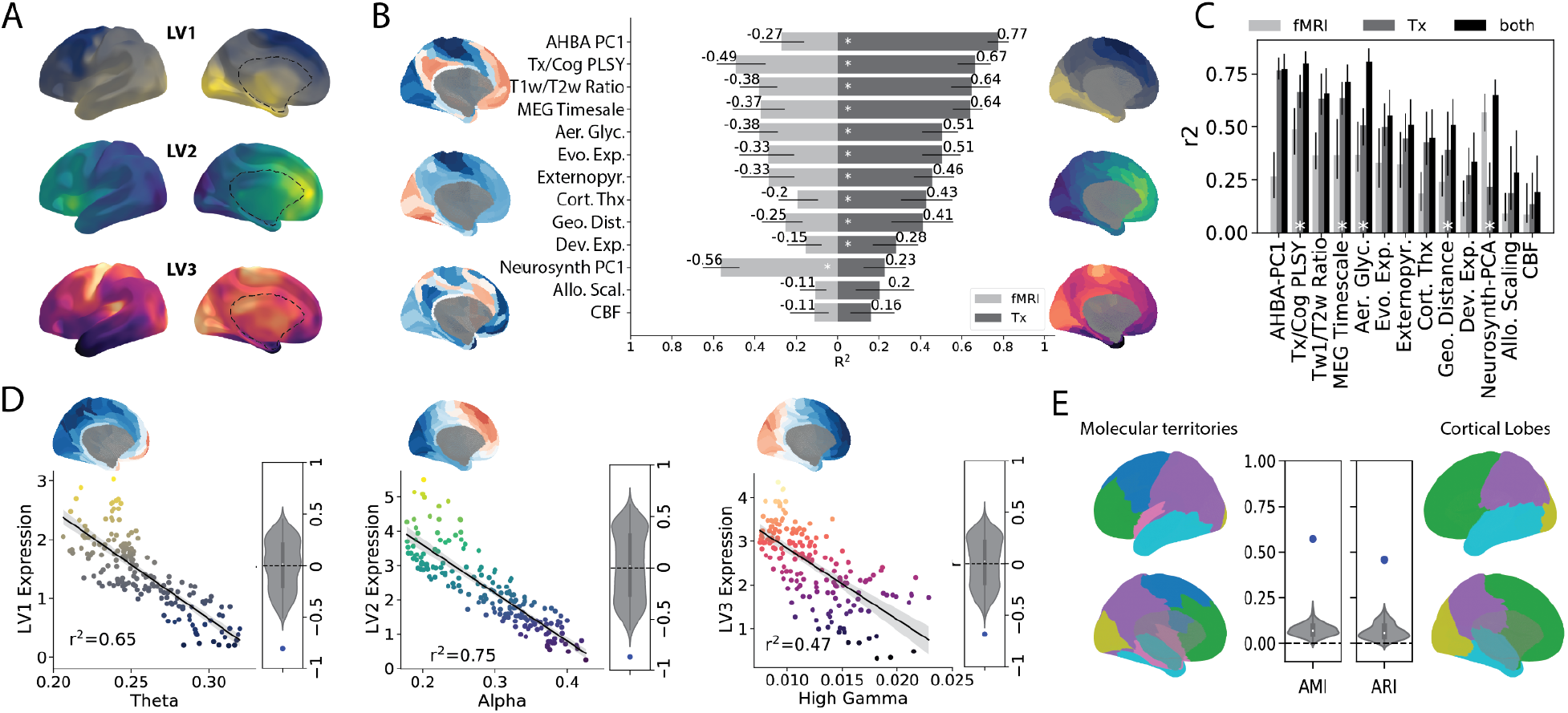
Cortical interaction of whole-brain transcriptomic gradients reflect fundamental properties of brain organization and function. Molecular gradients explain the topography of diverse brain properties and interact to form distinct functional territories. **A)** From top to bottom, a cortico-subcortical projection of PLS LV1, LV2 and LV3. **B)** PLS latent variables outperform fMRI gradients in predicting brain features. Each of the 3 PLS LVs, as well as the top three principal fMRI gradients from Margulies et al., 2016, were parcellated using the Glasser atlas. For several brain features (see Methods, **Table S3**), a linear model was fitted with the feature as the dependent variable and either the PLS LV (red) or fMRI principal gradient values (blue) as independent variables. Bars indicate the total model R^2^ (explained variance). Confidence intervals were obtained using bootstrapping. White stars indicate one model R^2^ was significantly larger than the other using bootstrap tests. **C)** Linear models were fit for the same features in B) with the feature as the dependent variable and either the three spatiomolecular axis values (from PLS, blue), the three fMRI principal gradient values (red) or all six variables as predictors (teal). Bars indicate the total model R^2^ (explained variance). Confidence intervals were obtained using bootstrapping. White stars indicate variables for which models incorporating both feature sets explained significantly more variance than models with only one feature set. **D)** Maps of cortical activity at different wavelengths acquired from MEG demonstrate topographic correlation to molecular gradients. Spatiomolecular axes and MEG maps were parcellated with the Glasser atlas and compared. The strongest correlation for each gradient is shown. Each dot represents a region; brains next to each graph represent cortical renderings of the respective MEG map. Beside each plot, the observed *r*-value of the correlation is compared to a null distribution of correlations created between the MEG map and 1000 permutations of the gradient maps generated using a conservative permutation procedure that preserves the spatial covariance structure. **E)** Cortical vertices were clustered to identify six molecular territories of overlapping spatiomolecular gradient concentrations (see **Fig S4A,B**). These molecular territories (left) were compared to (right) a surface rendering of lobar regions of interest, representing the occipital (yellow), parietal (purple), frontal (green) and temporal (turquoise) lobes. Adjusted mutual information (AMI) and adjusted Rand index (ARI) were used to compare the two vertexwise cortical maps. The blue dot represents the value, while the null distributions were created by using these two metrics to compare the lobar map to 1000 spun versions of the molecular territories.

### Candidate functional brain phenomena underlying molecular gradients

The previous analyses revealed that spatiomolecular gradients are in part spatially differentiable from the canonical functional gradients of the cerebral cortex. Yet, given that the interaction of molecular gradients gives rise to functionally distinct territories, we expected to find some evidence for differences in modes of functional communication along molecular gradients. Given previous literature has described traveling waves of cortical activity propagating across the cerebral cortex (Davis et al., 2020; Roberts et al., 2019; Zhang et al., 2018), we were curious whether these oscillations aligned with spatiomolecular gradients. We correlated spatial patterns of intrinsic spectral content of mass neuronal activity across several different wavelengths (measured using magnetoencephalography; MEG) with the three PLS latent variables. We found that each component strongly matched the spatial oscillation pattern of a different wavelength (**Fig 2D, S5**). LV1 matched theta activity (r = −0.81, p_spin_<0.001), LV2 matched alpha activity (r = −0.87, p_spin_<0.001) and LV3 best matched high gamma activity (r = −0.69, p_spin_<0.001). These results suggest that major components of subsecond neuronal communication occur preferentially along paths defined by spatiomolecular gradients, and may indicate a relationship between these two phenomena.

### Spatiomolecular gradients are differentially conserved across phylogeny

Previous analyses showed remarkable consistency in the expression of specific spatiomolecular gradients across several different datasets. Given extensive prior evidence linking spatial variation in expression across species and human development (Li et al., 2018; Zhu et al., 2018), we sought to evaluate whether spatiomolecular gradients were emergent (uniquely human) features of the brain, or whether they represent phylogenetically conserved features observable across the brains of other primate species. To evaluate this, we evaluated the expression of the three LVs using tissue samples extracted from adult macaque, chimpanzee, and a separate set of human brains (**Fig 3A**). Using the same approach described above (see **Fig 1I**), we tested how well the individual regional expression pattern of each LV matched the regional expression pattern of the discovery LVs compared to null models (**Fig 3A**). We found regional gradient expression to replicate well across humans (LV1 r=0.90-0.98; LV2 r=0.81-0.92; LV3 r=0.71-0.85), chimpanzees (LV1 r=0.92-0.96; LV2 r=0.80-0.93; LV3 r=0.46-0.72) and macaques (LV1 r=0.91-0.94; LV2 r=0.85-0.93; LV3 r=0.60-0.77). These individual correlations with the discovery dataset exceeded chance (p<0.05) for all individuals, except for but one chimpanzee individual for LV3 (**Fig S6**). A two-way ANOVA showed a significant species by component interaction (F[4;39]=3.43, p=0.017), where LV3 similarity was higher in humans compared to other primates. This suggests that LV3–which notably was the most variable across the cortex (**Fig S3A**)–is less conserved across primate species compared to LV1 and LV2.

We next tested whether the high similarity in spatiomolecular axes observed across human, chimpanzee and macaque brains was simply a product of generally conserved brain gene expression patterns across the three species, or whether conserved expression was especially enhanced for our three spatiomolecular axes. In other words, we wished to deduce how reproducible or conserved these gradients are compared to other spatial components of biologically covarying genes. Therefore, we derived 100 components of brain regional gene coexpression in the discovery dataset using principal components analysis. We used the interspecies reproducibility of these 100 components as a distribution against which to compare the expression pattern of the three spatiomolecular gradients, and we performed this analysis separately across each individual human, macaque and chimpanzee individual (**Fig 3B)**. Across all individuals, we found that reproducibility of LV1 and LV2 to be significantly (p<0.05) greater than expected for gene coexpression networks across all individuals. Across both species, LV1 and LV2 expression were consistently in the top 5% of biological components in terms of reproducibility. LV3 was generally less reproducible, though reproducibility was still within the top 10% of biological components for all except one chimpanzee individual. In all, we found strong evidence for the existence of homologous LV1 and LV2 gradients in chimpanzees and macaques, indicating conservation of these organizational phenomena at least across primate species.

**Figure 3:**
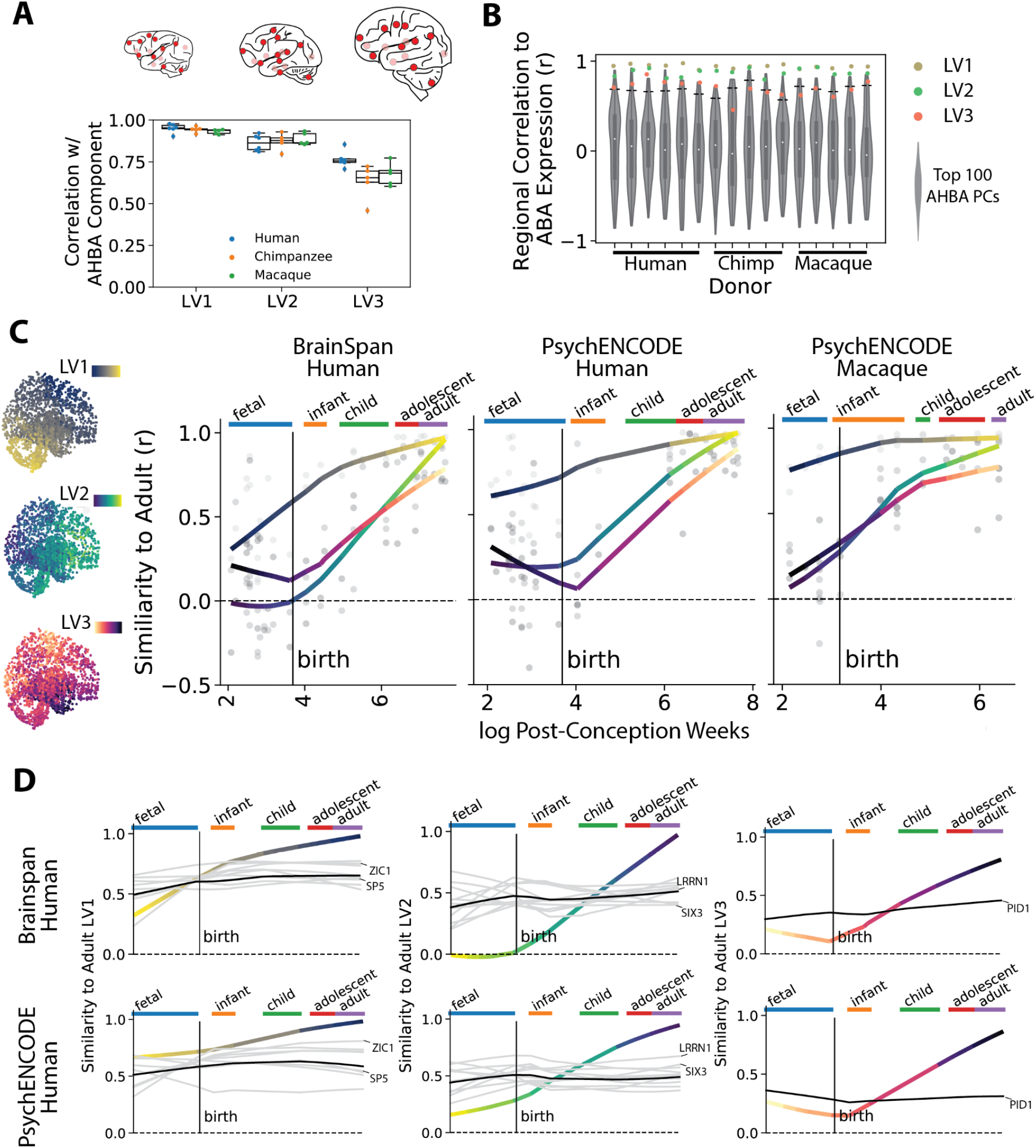
Tracking spatiomolecular gradients across phylogeny and ontogeny. The PLS latent variables are replicable across independent human datasets and primate species, and show distinct lifespan developmental trajectories. **A)** Six adult human, five adult chimpanzee and five adult macaque brains were available from the PsychENCODE dataset with tissue samples extracted from the same 16 brain regions (indicated with red dots). Component correlations (as described in Fig 1I) for each component were generated for each individual (see Figure S6 for individual null models). There was a significant component by species interaction, where LV3 expression was greater in humans than in other primates. **B)** To test whether high correlations were specific to the PLS latent variables over other sets of spatially co-expressed genes, for each individual we performed correlations described in A) for each of the first 100 principal components of AHBA gene expression data. Null distributions (gray) represent distribution of cross-dataset correlations. Horizontal lines indicate correlations that are in the top 5% of AHBA PC distribution. **C)** The method described in Fig 2A was applied to data from individual brain donors across the development spectrum in the BrainSpan dataset (left) and PsychENCODE human dataset (center) and PsychENCODE macaque dataset (right). Plots represent the relationship between log Age (in post conception weeks) and individual-level component correlations with AHBA (adult) PLS latent variables. Developmental gradient expression is remarkably similar across datasets, with adult-like LV1 expression present early in development, and adult-like LV2 and LV3 expression accelerating after birth. **D)** Clustering was used to identify a set of individual genes that demonstrate high regional expression similarity to adult LVs throughout brain development (i.e. are non-transitional). For all three LVs (columns) and both human datasets (rows), the y-axis represents regional similarity to adult AHBA LV expression. The trajectories of individual genes falling into the non-transitional clusters across both datasets are visualized juxtaposed against the trajectory of LVs. Showing high similarity to adult LV expression during prenatal development and maintaining this expression level into adulthood, these genes are candidates for establishment and maintenance of the LVs. A few example genes are highlighted to demonstrate consistency in gene trajectories across datasets. All genes are listed in **Fig S8**.

### Differential maturation of gradients observable across species

The directional radiation of the three spatiomolecular gradients resembles the morphogen diffusion patterns of whole-brain developmental spatial gradients (see **Fig 1A**). Therefore, we wished to assess whether the molecular gradients discovered in our adult samples are present during fetal brain development, and how they develop over the lifespan. We used expression patterns of genes shared across datasets to examine the development of the PLS LVs across two developing human datasets spanning post-conception week 8 to postnatal year 40, and one developing macaque dataset spanning post-conception week 8 to postnatal year 11. Specifically, we compared each individual’s PLS LV expression to that of the adult discovery set using regional correlations (from **Fig 1I).** Expression of adult-like gradients varied systematically with age in a gradient-dependent fashion (**Fig 3C, S7A, S8**). Supporting previous findings (Li et al., 2018), across all components, variance in expression was highest prenatally, with a few early prenatal individuals showing adult-like component expression across all three components (**Fig S7A)**. The developmental trajectories of gradient expression were remarkably similar across datasets and across species. Across all three datasets, correlation with adult expression of the LV1 gradient was already non-zero at early measurement, and slowly became more adult-like with age. In contrast, LV2 and LV3 expression was not observable at birth, but increased sharply throughout development, finally reaching adult-like levels at adulthood. However, a subset of LV-associated genes (see below, **Fig 4A**) showed an adult-like regional expression pattern throughout brain development, including during prenatal periods, across both human datasets (**Fig 3D, S7B, S8**) This set included genes known to be involved in areal patterning (e.g. SIX3, SP5, ZIC1). Most of these genes showed a similar developmental pattern in macaques as well, with a few notable exceptions (**Fig S8**).

### Molecular gradients are composed of genes relating to neural development and disease

Having established directional molecular gradients in the adult human brain, we were interested in understanding the roles of genes that contributed to them. We specifically focused on genes within the top 5% (adjusted for multiple comparisons, see Methods) of each component (**Fig 4A**). We found a clear and robust enrichment of terms relating to neural and neuronal development and morphogenesis, as well as synaptic signaling and extra-cellular matrix development (**Fig 4D, Table S4**). Furthermore, of 38 transcription factors that are known to be expressed in a gradient-like pattern in the developing mouse ventricular zone (Ypsilanti et al., 2021), 26% were included among our list of human gradient-associated genes (random gene set null mean = 3.3% [95% CI: 0.0%-7.9%], p_[perm]_ = 0.001). This included some well characterized morphogens and transcription factors (e.g., *PAX6, SIX3*), as well as genes known to exhibit regionally-specific or gradient-like expression in early development (*BCL11B, EPHA3, EPHA5, CDH6, NR2F2, PCDH8, SLN, TSHZ2*) and several members of the Wnt and cadherin/protocadherin family (**Table S1**). Interestingly, we also observed that multiple genes involved in gradient organization are known to be associated with neuropsychiatric disorders such as schizophrenia, autism, and bipolar disorder, demonstrating significant enrichment (**Fig 4E)**. To confirm these associations, we assessed significant enrichment of genes from each gradient in a multivariate GWAS across psychiatric disorders (Mallard et al., 2022). Genes in LV1 (p=0.026) and LV3 (p=0.0091) were enriched in factor one of this GWAS, which was associated with psychosis, mania, and depressive disorders. No enrichment was found for the second factor, which was composed of schizophrenia and bipolar I disorders (**Fig 4F**).

**Figure 4:**
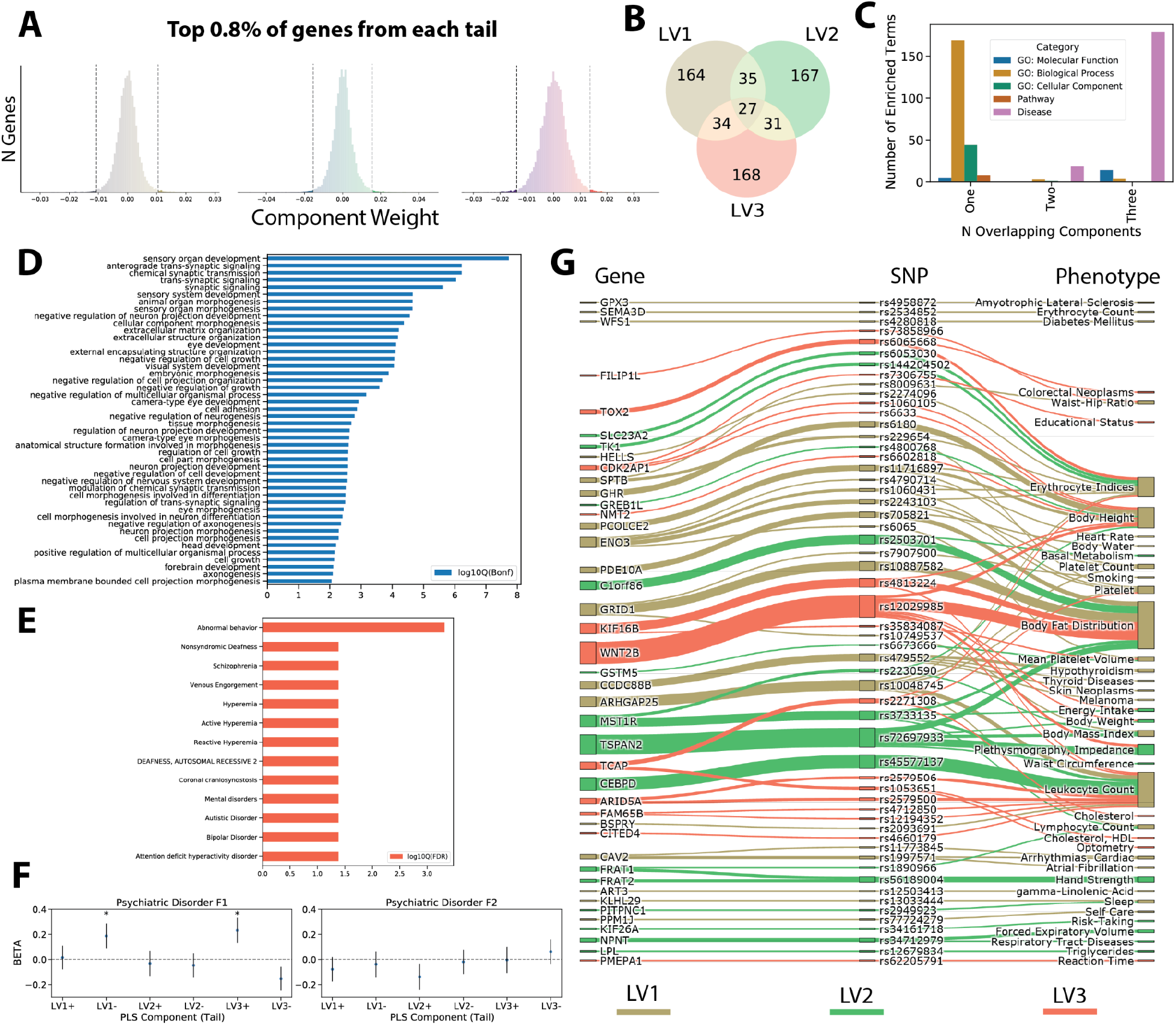
Annotation of gradient-associated genes. Genes composing adult LVs are associated with brain development, adult disease, and physiology. **A)** Distributions of PLS LV weights across genes. Genes weighted in the top 0.8% (5% / two tails / three components) of either tail of this distribution (130 per tail) were selected for annotation. **B)** Venn diagram showing how many genes were unique to one LV or shared among multiple LVs. **C)** Gene set enrichment was executed on genes that were involved in one, two or all three PLS latent variables. Genes belonging to only one component were mostly enriched for biological processes, whereas genes belonging to all three components were mostly enriched for diseases. **D)** Gene set enrichment across all gradient-related genes indicated a robust enrichment for terms associated with development and morphogenesis of neural organs and cells, as well as synaptic signaling. **E)** Genes contributing to molecular gradients were also enriched for several diseases, particularly psychiatric diseases. **F)** To garner greater specificity of the effects in E), gene sets from each tail were compared to a two-dimensional GWAS of neuropsychiatric disorders. Significant associations were seen between the first dimension and both LV1 and LV3 genes. The y-axis shows the effect size (standardized beta) of the association. **G)** We identified SNPs associated with brain expression of gradient-associated genes, and found genotype significant relationships those SNPs had with various phenotypes. Sankey diagram indicates these dependent relationships, suggesting pleiotropic effects of these genes on aspects of body morphology and circulation, as well as brain-related traits.

We next examined whether any overlap existed in the genetic anatomy of each LV. Despite the fact that the PLS approach enforces orthogonality among its LVs, we found that several genes contributed to more than one gradient, and 27 genes contributed to all three (**Fig 4B**). Interestingly, genes that contributed to only one gradient tended to be associated with specific biological processes but not diseases, whereas genes involved in all three components were associated with diverse neurological, neuropsychiatric and non-neural diseases but with few biological processes (**Fig 4C, Table S5**). Three of the genes highlighted earlier as candidates for early gradient establishment (*HRH3, TLL1, MGAT4C*) were also present in this set of multi-gradient, disease associated genes.

Finally, in order to gain a better understanding of direct relationships between brain gene expression and a broader array of relevant phenotypes, we leveraged a recently published phenotype-wide association study (PheWAS) using brain expression-based multi-ancestry meta-analytic quantitative association loci (mm-QTLs) (Zeng et al., 2022). Specifically, we identified single nucleotide polymorphisms (SNPs) that were significantly associated with expression of LV-associated genes in the brain, and which also showed genome-wide significant associations with biological traits and brain-related phenotypes (**Fig 4G, Table S6**). Many SNPs associated with brain expression of gradient genes were also associated with body size and distribution, metabolism and circulation. However, additional brain-related associations were noted, including amyotrophic lateral sclerosis (ALS) in LV1, sleep and risk-taking behavior in LV2, and educational attainment and cognitive traits in LV3.

## Discussion

Directional morphogenetic gradients are a well-established organizational element of the developing brain, serving as scaffolds for the coordination of functional arealization. We demonstrate that these gradients are retained in the adult human brain, in the form of robust and systematic transcriptional variation along three spatially embedded brain axes. These spatiomolecular gradients traverse the cerebral cortex along paths of intrinsic neural oscillations, and they interact to delineate territories representing well described anatomical and functional cortical divisions. In all, we describe a reproducible pattern of brain gene expression that resembles developmental morphogen gradients, and which interacts in the cerebral cortex to support functional differentiation in the adult brain. Coordinated prenatally through morphogen diffusion and consolidated throughout development, these spatiomolecular gradients likely serve as fundamental organizational axes of brain functional organization.

Functional gradients in the adult human cerebral cortex have been a topic of concerted study in recent years, where a sensorimotor-association axis has been proposed as a major feature of functional organization that can also explain many aspects of cortical physiology and neurobiology (Margulies et al., 2016; Sydnor et al., 2021). In parallel, the introduction of transcriptomic datasets with dense cortical sampling have allowed the establishment of intrinsic components of cortical gene expression (Burt et al., 2018; Hawrylycz et al., 2015; Markello et al., 2021; Wagstyl et al., 2022). Several studies have further shown a spatial correspondence between these principal axes of function and gene expression (Fulcher et al., 2019; Hansen et al., 2021; Oldham et al.; Wang et al., 2022). Our approach differed from that of previous studies in two critical ways: i) we used data from the entire brain rather than only the cerebral cortex; and ii) we enforced a degree of supervision by constraining transcriptomic covariation to directional radiation, in accordance with the distribution of theoretical developmental gradients (which are embedded in Euclidean space). The resulting transcriptomic gradients bore some spatial resemblance to fMRI-based functional gradients, but were far more similar to patterns of neural oscillations measured at millisecond timescales. These findings corroborate previous studies finding strong relationships between cortical gene expression and millisecond timescale functional activity (Gao et al., 2020). The directional pattern of our molecular gradients was also reminiscent of genetic variation in cortical morphology (Chen et al., 2012, 2013; Valk et al., 2020). In fact, the molecular gradients explained significantly more variance than functional gradients in the spatial distribution of most physiological cortical features tested. In contrast, the fMRI-based functional hierarchy map (Margulies et al., 2016) far better explained the topography of meta-analytic task-related coactivation patterns from the Neurosynth database. This is notable given that electrophysiological and fMRI measure brain activity at different time-scales, overlapping mostly at the high frequency band (Baria et al., 2011; Hermes et al., 2017; Kucyi et al., 2018; Leong et al., 2016; Shafiei et al., 2021). Together, these results suggest that our spatiomolecular gradients and previously described functional gradients probably represent two distinct elements of cortical organization. As such, we hypothesize that spatiomolecular gradients dynamically coordinate physiological properties of the brain, and their confluent cortical topographies enforce the distribution of functionally distinct regions in space. In contrast, we speculate that the primary functional gradients help to coordinate optimal communication between these spatially distinct regions, forming networks of long-range projections that likely manifest as functional networks measurable with fMRI (Achard and Bullmore, 2007; Bajada et al., 2019; Chen et al., 2017; Sepulcre et al., 2010; Wang et al., 2022). Both organizational elements are likely reflected by independent functional activity at different timescales (Kiebel et al., 2008; Murray et al., 2014), a notion supported by our finding that functional and spatiomolecular gradients each contributed additive information in explaining the spatial distribution of brain energy consumption (aerobic glycolysis).

The role of morphogen gradients in the coordination of downstream molecular events during neurodevelopment is well established (Cadwell et al., 2019; Hébert and Fishell, 2008). We show that these molecular trails are represented in the adult brain by patterns of genes coexpressed along similar axes. The genes composing these molecular axes included many known developmental morphogens and transcription factors, and were enriched almost exclusively for biological processes relating to neural organ and neural cell development. On the other hand, the molecular axes were not reliably observed in prenatal brains, and instead only became maximally recognizable in adulthood, potentially due to differential contributions from functionally-related genes. Previous literature has described waves of transcriptomic changes that occur along stereotyped developmental schedules (Habib et al., 2016; Li et al., 2018; Telley et al., 2019). As tissue samples used to discover these molecular gradients were sourced exclusively from adult donors, the resulting gradients likely represent a confluence of multiple waves of transcriptional change consolidating the axes that begin to be established during early development – a so-called “palimpsest” model as described previously (Hallgrímsson et al., 2009). This idea is supported by our finding that a subset of the top gradient-associated genes for LV1 and LV2 demonstrated gradient-like expression patterns prenatally and throughout development. Some of these genes may be involved in the early establishment and/or long-term maintenance of spatiomolecular gradients, which are otherwise consolidated across development by the recruitment of many other genes with differing functional consequences.

The spatial axes along which these gradients vary are phylogenetically ancient. Rostral-caudal and dorsal-ventral identity are evolutionarily conserved aspects of body-plan formation that are genetically encoded (Brooun et al., 2022; Meinhardt, 2008; Petersen and Reddien, 2009). Many similar signaling pathways also encode these spatial axes in the brain to create explicit divisions (Meinhardt, 2008). Some examples of this organizational feature include the fact that phylogenetically ancient species exhibit explicit structural divisions separating brain segments along the rostral-caudal axis (Dutel et al., 2019), that cell lineage studies find biased allocation along rostral-caudal brain axes in early development (Fasching et al., 2021), and that cerebral organoids spontaneously self-organize around spatial axes (Kadoshima et al., 2013). Therefore, it is unsurprising to find excellent reproducibility of the human-derived rostral-caudal (LV1) and dorsal-ventral (LV2) spatiomolecular axes in various non-human primate species. In contrast, LV3 showed the most variation within the cerebral cortex, a brain region of disproportionate size in primates, and in humans compared to other primates (Hofman, 2014). Interestingly, this gradient was not as reproducible in non-human primates, was made up of genes enriched for human neuropsychiatric disorders and associated with educational attainment and cognition, and was expressed in a pattern similar to gamma waves observed during complex cognition (Herrmann et al., 2010). This component also continued to consolidate throughout childhood and adolescence and demonstrated few genes with adult-like expression during early development, as opposed to LV1, which showed a fairly adult-like distribution already during early life. Together, these findings outline how various molecular gradients interact during different epochs of brain growth to guide regional differentiation and functional specialization.

Previous studies suggest that cortical expansion can allow greater cortical differentiation, particularly along spatial axes (Imam and Finlay) similar to those characterized by our PLS analysis (Hill et al., 2010; Reardon et al., 2018). Therefore, a gradient organizational framework can create an intimate relationship between (brain) compartment size or shape and the complexity of regional differentiation (Lau et al., 2021). This may be an important principle underlying the observation that specialized function co-occurs with nonlinear variation in brain morphology, even among conspecifics (Hecht et al., 2019). This also may explain the finding that LV3 was less reproducible in other primate species with less expanded cortex. However, the directionality of the relationship between brain shape/size and gradient expression is still not clear. We found that many of the gradient-associated genes in this study showed pleiotropic effects on human physical form and function. This finding supports the observation that genes regulating brain axes may play a similar role on the axes of other organs. In addition, in a scenario where organ size relates to expression of gradient-forming genes, co-expression of genes supporting the attainment and/or maintenance of larger size may also be expected. This may explain the observation that mutations within some of the LV-associated genes in our study are linked to interindividual variation in metabolic and anthropometric traits.

Our analyses suggest that spatiomolecular gradients are highly conserved across adult individuals, underscoring the importance of both the establishment and maintenance of these gradients for healthy brain functioning. Previous literature suggests disruption of the formation of early developmental gradients through mutations to gradient-associated genes can cause mortality and severe developmental disorders (Flores-Sarnat and Sarnat, 2007). Our analysis supports this premise, as a small set of genes contributing to all three adult brain gradients were strongly enriched for disease associations. These genes may play an important role in anchoring one or more gradients, and their disruption could impact morbidity through complex perturbation across multiple components of brain organization. We furthermore found that genes associated with adult LVs are enriched for several psychiatric disorders and demonstrated overlap with a multifactorial psychiatric disease GWAS (Mallard et al., 2022). Though the causal relationship between brain disease and the establishment or consolidation of molecular gradients is currently unknown, their association further underscores that molecular gradients are likely to be important elements in the emergence of brain-related traits.

There are several potential limitations that must be acknowledged while interpreting the present findings – the foremost of which is the reliance on whole-brain spatial correlations to draw inference. While the comparison of densely sampled, multi-modal whole-brain information is a distinct strength of our approach, correlations of the spatial topography of two signals, no matter how high they may be, are not indicative of complex relationships. We tried to mitigate this limitation by using analysis-specific permutation tests and spatially-aware null models to provide conservative testing for significant relationships. Still, there are no experimental manipulations in this study and we therefore cannot establish causality. Many state-of-the-art null modeling approaches in neuroimaging seek to correct for spatial autocorrelation (Markello and Misic), whereas this was the phenotype of interest in the present study. We overcame this limitation by validating our molecular gradients in several independent datasets, albeit these datasets had much more limited spatial sampling of the brain. Bioinformatic annotation of gene sets also comes with numerous limitations. Transcriptomic information does not represent protein concentration with great fidelity (Schwanhäusser et al., 2011), and relationships between gene sets and enriched terms does not entail a causal relationship between our gradients and biological processes or diseases. We addressed this limitation by validating disease associations with different techniques, and by identifying phenotypes relating to SNPs that were themselves associated with brain gene expression of LV-associated genes. However, each of these approaches is not without its own set of limitations. Finally, we attempt to further protect against other limitations of data analysis by publishing all code used to analyze the present data in order to maximize reproducibility.

This study establishes directional gene expression gradients as measurable phenomena in the adult human brain and key features of functional organization in the cerebral cortex. While many contemporary studies consider autocorrelated gene expression information an artifact or nuisance to be statistically removed (Wei et al., 2022), the role of this kind of signal must be acknowledged as a biological phenomenon with an important role in brain organization. Indeed, gene expression gradients are present across many organs (Gilmour et al., 2017; Schwank and Basler, 2010), likely due to their capability of optimizing tissue differentiation and enforcing organizational features such as symmetry and functional distribution. Future work should strive to characterize the developmental sequences of these gradients with greater temporal granularity, and further demonstrate how variations in this process influence brain and behavior at the individual level.

## Acknowledgements

The authors would like to acknowledge the various open data initiatives that provided all data used in this manuscript, and therefore made this work possible. In addition, we would like to thank Casey Paquola for input and feedback on this manuscript. JWV (T32MH019112), JS (T32MH019112) and AP (F31 MH123063-01A1) would like to acknowledge funding from the NIH. KSW was supported by the Wellcome Trust (215901/Z/19/Z). AAB and JS were supported by K08MH120564. JYH acknowledges support from the Helmholtz International BigBrain Analytics & Learning Laboratory and the Natural Sciences and Engineering Research Council of Canada. VJS was supported by a National Science Foundation Graduate Research Fellowship (DGE-1845298). BB acknowledges research support from the National Science and Engineering Research Council of Canada (NSERC Discovery-1304413), the Canadian Institutes of Health Research (FDN-154298, PJT-174995), SickKids Foundation (NI17-039), Azrieli Center for Autism Research (ACAR-TACC), BrainCanada (Future Leaders), FRQ-S, and the Tier-2 Canada Research Chairs program. BB was furthermore funded in part by Helmholtz Association’s Initiative and Networking Fund under the Helmholtz International Lab grant agreement InterLabs-0015, and the Canada First Research Excellence Fund (CFREF Competition 2, 2015-2016) awarded to the Healthy Brains, Healthy Lives initiative at McGill University, through the Helmholtz International BigBrain Analytics and Learning Laboratory (HIBALL).

## Methods

All statistical analyses were performed and plots generated using the python programming language (v. 3.7.3), mainly using the numpy, scipy, pandas, sklearn, statsmodels and seaborn libraries. Code and library versions necessary to reproduce all analyses described can be found at https://github.com/PennLINC/Vogel_PLS_Tx-Space.

### Samples and Preprocessing

#### Discovery Sample

The genomic gradients described and analyzed throughout the present study were generated using data from the Allen Human Brain Atlas. A detailed description of this dataset can be found elsewhere (Hawrylycz et al., 2012). Briefly, the dataset consists of 3702 tissue samples extracted from eight cerebral hemispheres across six human donors (1 female, ages 24.0-57.0, mean=42.50 +/− 13.38). Each tissue sample underwent high throughput microarray mRNA analysis across 58,692 probes and underwent preprocessing as previously described (Arnatkeviciute et al., 2019; Markello et al., 2021). The 58,692 x 3702 data matrix downloaded from (https://human.brain-map.org/static/download) was further processed using the abagen toolbox (version 0.1.3; https://github.com/rmarkello/abagen) (Markello et al., 2021) based on prior recommendations (Arnatkeviciute et al., 2019). Rather than averaging tissue samples within regions of a brain atlas, all samples were extracted and underwent preprocessing. The following text was generated directly from abagen, describing the exact methodologies employed during preprocessing:

First, microarray probes were reannotated using data provided by (Arnatkeviciute et al., 2019); probes not matched to a valid Entrez ID were discarded. Next, probes were filtered based on their expression intensity relative to background noise (Hawrylycz et al., 2015), such that probes with intensity less than the background in >=50% of samples across donors were discarded. When multiple probes indexed the expression of the same gene, we selected and used the probe with the most consistent pattern of regional variation across donors (i.e., differential stability; (Hawrylycz et al., 2015). The MNI coordinates of tissue samples were updated to those generated via non-linear registration using the Advanced Normalization Tools (ANTs; https://github.com/chrisfilo/alleninf). Inter-subject variation was addressed by normalizing tissue sample expression values across genes using a robust sigmoid function (Fulcher et al., 2013). Normalized expression values were then rescaled to the unit interval. Finally, gene expression values were normalized across tissue samples using an identical procedure. Samples assigned to the same brain region were averaged separately for each donor and then across donors, yielding a regional expression matrix. After processing, a final 3466 sample x 15,634 gene matrix was used for subsequent analysis.

#### Replication cohorts

Several publicly accessible datasets were used to replicate and extend results. The Brainspan “Developmental Transcriptome” dataset consists of 524 cortical, subcortical and cerebellar tissue samples extracted from the brains of 42 donors (19 female, ages 8 post-conception weeks to 40 years, mean=86 +/− 137 post-conception months). Each tissue sample underwent mRNA sequencing for 47,808 unique genes, and were preprocessed using methods previously described (Miller et al., 2014). This data was downloaded from https://www.brainspan.org/static/download.html in 2020. After removing duplicates, 13,750 genes were present in both the Allen Brain Atlas discovery dataset and the present Brainspan dataset, and this 524 sample x 13,750 gene dataset was used for analysis. To maximize consistency with the discovery datasets, initial reproducibility analyses were conducted only among the six adult donors in the Brainspan dataset (3 female, age 21-40, mean = 31.2 +/− 7.2).

A total of 2,483 cortical, subcortical, midbrain and cerebellar samples were extracted from the brains of 376 donors (105 females, age 20-75, mean = 58.7 +/− 9.7) from the NIH Genotype-Tissue Expression (GTEx) dataset Version 8(GTEx Consortium, 2013). Each sample underwent mRNA sequencing for 15,758 unique genes. In order to obtain individual (rather than binned) ages, protected GTEx data was downloaded under accession number 26317. However, all other GTEx data besides age can be downloaded from (https://gtexportal.org/home/datasets). 12,647 genes overlapped with the discovery sample, and only data for these genes were used for subsequent analysis. After principal component analysis across technical variables, the first five principal components (PCs) were regressed from the expression data of each gene by finding the residual of an ordinary least squares regression model with expression data as the dependent variable and the five PCs as predictors. Previous work with the GTEx dataset describes removal of subjects based on presence of brain diseases and other potentially confounding factors (Hartl et al., 2021). To rule these factors out as drivers of our results, we reproduced our results after subsampling the original 376 donors to only include 227 individuals included in (Hartl et al., 2021), known to be free of brain diseases. Furthermore, to better match the age range of the initial discovery sample and to rule out advanced age as a drive of the results, initial reproducibility analyses were also repeated in both the original (n=178) and the subsampled (n=121) GTEx datasets after excluding individuals above the age of 60.

To assess whether our results generalized across species, we leveraged a pre-curated dataset described in (Zhu et al., 2018). This dataset combines tissue samples from the six adult humans, five adult chimpanzee brains and five adult macaque brains. The dataset can be downloaded from http://evolution.psychencode.org/#, where it is labeled as “Adult human, chimpanzee, macaque data” in the mRNA-seq tab. The dataset is pre-harmonized to include a consistent set of 16 cortical and subcortical brain regions with mRNA sequencing performed on 11,346 sets of carefully curated homologous genes (Zhu et al., 2018). All but five of these genes were also present in the Allen Brain Atlas discovery dataset, and only the missing five were excluded for subsequent analysis. Similarly, to replicate developmental findings across species, we used a second pre-curated dataset, downloadable from the same link, labeled “Developmental rhesus and human data”. This dataset, described in (Zhu et al., 2018), includes tissue samples from 36 human brains (15 female, ages 8 post-conception weeks to 40 years, mean = 97 +/− 147 post-conception months) and 26 macaque brains (8 female, ages 60 post-conception days to 11 years, mean = 36 +/− 46 post-conception months). The dataset includes brain regions extracted from 16 cortical, subcortical and cerebellar brain regions across both species. Three transient developmental brain regions were excluded (lateral, medial and caudal ganglionic eminence), while other prenatal regions were considered equivalent to their most similar adult brain regions (e.g. dorsal thalamus to mediodorsal thalamus, upper rhombic lip to cerebellum, etc). This was only relevant for two of 62 total brains that possessed these early developmental regions. Each brain region had mRNA sequencing performed on 27,932 genes, of which 13,113 overlapped with the Allen Human Brain Atlas discovery set and were used for analysis.

### PLS Methods

#### Model fitting, optimization and validation

Observations in developing brains suggest the presence of multiple large-scale directional gene expression gradients that exist within and across compartments (**Fig 1A**), and which help to coordinate developmental programming and functional specialization. The current study investigated whether such gradients are present in the adult brain. Our objective was therefore to identify patterns of autocorrelated brain gene expression that vary systematically along a 3-dimensional linear axis. To accomplish this goal, we used a cross-decomposition framework in order to determine modes of latent covariance between gene expression (**X**; a 3466 sample by 15,634 gene matrix) and euclidean space (**Y**; a 3466 sample by 3 dimension matrix). Under a prediction framework, such a model would use latent gene expression components to predict the 3-dimensional location (i.e. x-,y- and z- coordinates) of a tissue sample in space (**Fig S1A**). However, the gene expression component (PLS X) in this case will be a latent variable representing linear variation in gene expression across a 3-dimensional axis – a construct consistent with the transcriptomic gradients we wish to characterize. Note here that, due to left-right symmetry relative to other brain dimensions, absolute x-coordinates were used in model fitting. This is equivalent to projecting all tissue samples onto one hemisphere, but otherwise maintaining their distance from the origin and y- and z-coordinate. True x-coordinates were used for display purposes only.

As a first step, principal components analysis (PCA) was used to decompose the gene expression data into its first 100 components. These 100 components collectively explained 84.6% of the original data and reduced the input matrix X to a 3466 sample by 100 component matrix (**Fig S1A**). The choice to precede cross-decomposition with this data reduction step was made for the following reasons: i) recent benchmarking work suggests the stability of cross-decomposition weights is greatly reduced as a function of feature/sample ratio, such that a smaller feature/sample ratio is desirable (Helmer et al., 2021); ii) gene expression data exhibits a hierarchical covariance structure with biologically meaningful co-expression networks at various scales (Hawrylycz et al., 2015); iii) computational efficiency is greatly increased, allowing for rapid permutation and ease of reproducibility.

Next, a 70/30 train-test split was applied to the dataset. The Allen Brain Atlas dataset provides 212 structural labels indicating the exact brain region the anatomist extracted the tissue from. The train/test split was stratified such that each of these tissue samples was equally distributed across the training and test sets. Across splits, we ensured components were inverted so as to match the direction of the full sample output. The following optimization procedures were executed exclusively on the n = 2,599 sample training set. A miniature grid search assessed model performance, estimated using both R^2^ score and mean squared error, varying across different estimators (partial least squares regression [PLSR], partial least squares canonical [PLSC] and canonical correlation analysis [CCA]) and number of components (1,2, 3; note that 3 is the y matrix rank and therefore the maximum allowable number of components). For each of these nine parameter sets, out-of-sample performance was obtained using ten sets of ten-fold cross validation. This analysis revealed a three-component PLSR model to provide the best out-of-sample performance (**Fig S1C**), which was used for subsequent analysis. For completeness, this process was repeated without the initial PCA data reduction step, but this resulted in poorer model performance.

To ensure each component predicted unique variance greater than chance given the data structure, the 3-component PLSR was fit 1000 times on permuted data (i.e. with shuffled labels) to create a null model distribution (**Fig S1D**). Given the spatially autocorrelated nature of gene expression data, it is common to perform “spatially-aware” null models (Markello and Misic) that create null permutations with spatial autocorrelation consistent with the initial dataset. However, such an approach is inappropriate in this context given that autocorrelation is the target of the present analysis rather than a confounder to it. Instead, we validated the PLS LVs by reproducing them in several external datasets (see below).

We assessed the final model fit by using the 3-component PLSR model fit to the entire training set and using it to predict the x-, y- and z-coordinates of the n=867 sample left out test set. Once satisfied with the predictive performance of the model, we fit the model to the entire dataset and obtained predicted x-, y- and z- coordinates for all tissue samples. We also calculated the absolute error (predicted - actual) in mm of every sample along all three axes (x,y,z), as well as the mean of these three values. The mean error was correlated with a measure of extremity, representing the mean distance between a sample and every other sample. We also correlated error against distance from origin, measured as the distance of the sample to MNI coordinates [0,0,0].

We also performed a robustness test to ensure the PLS results were not somehow influenced by the intrinsic orientation of the brain (**Fig S1F**). In the training set, we performed 100 random rotations of the brain and applied these rotations to the y, z, and (absolute) x coordinates of each sample. We then conducted the same three-component PLS analysis between gene expression and Euclidean coordinates, though this time the Euclidean coordinates were randomly rotated. We compared empirical LVs rotated LVs by looking at correlations between their PLS X loadings. If the empirical (unrotated) spatiomoelcular gradients were robust, we would expect the latent Y (spatial) variables to change but the latent X (gene expression) variables to remain consistent. For each new rotated PLS model, we sequentially (from first to last rotated LV) assigned each LV to its most similar (remaining) empirical LV by finding the (remaining) empirical LV to which the rotated LV had the highest PLS X weight correlation. Once an empirical LV was matched, that LV was removed for the next LV in the sequence. In other words, if rotated empirical LV1 was most similar to empirical LV1, rotated LV2 was only compared to empirical LV2 and LV3. Once all rotated LVs were assigned, we created a correlation matrix of PLS X loadings of all 300 LVs (3 LVs x 100 rotations), and ordered them by their previously described empirical LV similarity. This showed that gene-expression LVs were highly similar across permutations despite spatial rotation. Furthermore, the rotated LVs were highly similar to the empirical LVs; the mean R^2^ representing shared variance between empirical rotated and X loadings was 0.91 for LV1,0.71 for LV2 and 0.78 for LV3 (**Fig S1F**).

#### Evaluation of PLS latent variables

Bootstrap resampling with replacement was used to generate confidence intervals around loadings for each of the three PLS latent variables (LVs). The confidence intervals were used to generate p-values representing the likelihood that the loading crosses 0, and these p-values were subsequently FDR corrected across loadings. To further characterize genes contributing to each component, loadings with FDR Q>0.05 were considered unstable and were set to 0. These newly regularized PLS component loadings were then transformed back into individual gene space by finding the dot product between the loading vector and the transposed principal component matrix, the latter of which was standardized before multiplication (**Fig S1B**).

To evaluate whether PLS LV expression varied meaningfully with aspects of brain developmental spatial organization, each tissue sample was categorized based on its anatomical label (**Table S7**). For visualization purposes, samples were given a broad spatial categorization into one of the following areas: Frontal, Temporal, Occipital, Parietal, Sensorimotor, Limbic, Subcortex, Brainstem, Cerebellum. In addition, samples were given a label based on their cortical type: association cortex, sensory cortex, cerebellar cortex, or not cortex. Samples were further categorized based on whether the region from which they were extracted originates from the myelencephalon, metencephalon, mesencephalon, diencephalon or telencephalon. Similarly, for all samples not categorized to the telencephalon, samples were categorized based on whether the region of extraction originates in the dorsal or ventral plate of the developing brain. Based on the hypothetical expression pattern of LV1 and LV2 (**Fig 1A**), LV1 expression was compared to the neural tube segment categorization, and LV2 to the dorsal/ventral plate categorization. Finally, the cortical expression of LV3 was compared to cortical T1w/T2w ratio (see section “**Comparison to canonical brain features”** below for details of surface representation of PLS LVs and T1w/T2w ratio).

#### Out-of-sample replication procedure

Several external datasets include similar gene sets to the Allen Human Brain Atlas, allowing for harmonization of the PLS X variables. In contrast, x-y-z coordinates were not available in external datasets, and spatial sampling was considerably sparser. However, given that the gradients defined in these analyses span the entire brain, we expected that the spatial distribution of samples in external datasets was wide enough to detect the presence of the gradients.

The replication procedure (**Fig 3A**) involved the following steps: First, PLS-derived gradients were downsampled to match the resolution of external datasets by finding the mean value (for each component) of all samples falling inside of regions sampled in external datasets (see **Table S7** for mapping between Allen samples and regions from external dataset). For example, for replication in the BrainSpan dataset, mean values were derived from within the frontal, temporal, occipital, and parietal lobes, sensorimotor cortex, striatum, thalamus and cerebellum. Importantly, correlations were never performed across any analysis described below (group or individual) when n regions < 6.

Next, PLS X gradients were derived in external samples. This involved refitting both the PCA and the PLS model in the Allen Human Brain Atlas discovery set using only genes that overlapped between both the discovery and replication sets. 10-fold cross-validation was employed in the discovery set to compare overall performance of this PLS model compared to the original model. Next, the newly fit PC model was used to transform the replication samples to achieve a sample by (gene expression) component matrix, and the newly fit PLS model was then applied to this matrix. Note that prior to this step, the replication dataset was MinMax transformed so that all values fell between 0 and 1, using samples most similar to the Allen Human Brain Atlas dataset age range as the reference sample. Specifically, adult samples were used as the reference for BrainSpan, “younger” healthy controls (see above) were used as the reference for GTEx, and adult humans were used as reference in the PsychEncode datasets. This entire procedure together resulted in expression of each PLS latent variable for each sample in the replication dataset.

Finally, replication strength was evaluated by finding the correlation between regional PLS X expression in the (downsampled) AHBA data and PLS X expression in the external dataset. In effect, this analysis determines the degree to which the genes associated with a PLS latent variable are expressed in a regionally consistent manner across both datasets. A higher correlation indicates regional gradient expression more similar to the AHBA dataset. Consistent expression indicates a similar gradient exists in the external dataset (insofar as brain regions share the same spatial distribution), and further allows analyses to be run in the external dataset with the assumption that the gradient at hand is consistent with that described in the discovery dataset. To evaluate how well the gradients generalized across datasets, this replication procedure was performed specifically on the mean of all adult human samples in three separate datasets: BrainSpan, GTEx and PsychEncode (see **Replication cohorts**). Importantly, this procedure can be applied to test the consistency of a full dataset to the Allen dataset (by averaging expression across donors), but can also be applied at the individual donor level.

### Cortical surface analysis and transcriptomic territories

#### Surface rendering

The PLS analysis was conducted on samples extracted from the entire brain, including cortex, subcortex, midbrain, cerebellum and brainstem. However, we were interested in assessing whether whole-brain gradients converge in any functionally meaningful way in the cerebral cortex, where transcriptomic variation is considerably less pronounced (Hawrylycz et al., 2012). For each PLS X component, a 7 mm cube was created centered around the MNI coordinates of each tissue sample, where the values corresponded to PLS X expression of that component. This set of cubes was saved as a nifti file and converted to a fslr32k left hemisphere surface using connectome workbench’s “volume-to-surface-mapping” command. Values were then interpolated across the cortical surface using the “metric-smoothing” with the “fix-zeros” flag and a 5 mm kernel, so that all vertices contained values for each gradient.

#### Derivation of canonical brain features

We were interested in whether gradient distribution could explain the topography of other canonical cortical features. Canonical features chosen were those used in Sydnor et al. (Sydnor et al., 2021): T1w/T2w ratio, evolutionary expansion, allometric scaling, cerebral bloodflow, Neurosynth PC1, externopyramidization, aerobic glycolysis, cortical thickness and Allen Human Brain Atlas PC1. To this list of features, we added maps for average geometric distance as described in Margulies et al. (Margulies et al., 2016), developmental cortical expansion, magnetoencephalography intrinsic timescale, and gene expression-associated cognitive functional activation as described in Hansen et al. (Hansen et al., 2021). **Table S3** describes each feature and the paper and dataset from which they are sourced. Values for each feature were available using the Glasser parcellation (Glasser et al., 2016).

#### Comparison to principal fMRI gradients

Recent work suggests the principal functional gradients, derived using data-driven decomposition of resting-state functional connectivity (Margulies et al., 2016; Vos de Wael et al., 2020), represent a fundamental cortical hierarchy that discriminates several functional, morphological and biological features of the brain (Sydnor et al., 2021). Given that the gradient territories derived in this analysis also discriminated brain features, we conducted a direct comparison between these organizational brain maps in predicting the distribution of different cortical features. To achieve this goal, for each of the brain features previously discussed, we fit three ordinary least squares linear models with brain features as dependent variables; one used the three molecular gradients from the present analysis as independent variables, one used the first three principal functional gradients (Margulies et al., 2016), and one used all six gradient maps. We recorded the total explained variance (R2) of each model. To test whether one model explained significantly more variance than another, we performed bootstrapping resampling (100 iterations with replacement) to generate confidence intervals. A significant difference was recorded when the lower 95% CI of the higher distribution did not overlap with the mean of the lower distribution.

#### Clustering analysis and molecular territories

Clustering analysis was used to find empirical and data-driven territories of gradient overlap. The cortical array of each PLS X component was concatenated creating a 32,492 vertex x 3 component matrix, which was normalized with a MinMax scaler so that all values fall between 0 and 1. This matrix was subjected to hierarchical agglomerative clustering using euclidean distance and ward criterion for linkage, varying number of clusters (k) between 2 and 15 clusters. Silhouette score (Rousseeuw, 1987) and Calinski-Harabasz score (Calinski and Harabasz, 1974) were used to evaluate the fit of each clustering solution, where “peak” solutions with better scores relative to surrounding solutions were sought. Both score types converged in demonstrating a peak at k=6 (**Fig 2B**). Labels from the 6-cluster solution were subsequently obtained for each vertex and visualized. For the purposes of interpretation, the distribution of each PLS X component was visualized for each cluster. To compare the molecular territories to canonical cortical lobes, a glasser-space parcellation was obtained that assigned each parcel to either the parietal, occipital, frontal or temporal lobe (Keller et al., 2022). This map of lobar labels was compared directly to the glasser-space parcellation of molecular territory labels using adjusted mutual information score and adjusted Rand index – two approaches to compare two clustering solutions. To compare these results to chance, we calculated the same scores for 1000 spatially-aware (i.e. spun) null permutations (see section **Comparison to electrophysiological brain activity** below) of the molecular territory map.

#### Neurosynth decoding

We performed neurosynth decoding to further establish relevance of transcriptomic territories to functional organization. The objective of this analysis was to ascertain behavior-relevant terms associated with meta-analytic functional activation of regions falling within each transcriptomic territory. The neurosynth v4-topics-100 association maps were downloaded from (https://neurosynth.org/analyses/topics/v4-topics-100/). Each of these 100 maps represents regional meta-analytic functional coactivation associated with a set of associated terms, derived using latent dirichlet analysis (see (Yarkoni et al., 2011) for details). Thirty-two maps were removed from the analysis because the associated terms were not associated with behavior (see **Table S8**), Each map was subsequently converted from volume to surface using the same approach as above (but without the interpolation step). For each topic map, the mean value was extracted within each transcriptomic territory. The top 5 topic maps for each territory were recorded and visualized.

#### Comparison to electrophysiological brain activity

Seven magnetoencephalography (MEG) cortical surface maps were downloaded using the Neuromaps software (Markello et al., 2022). These maps approximate local field potentials from cell populations by detecting electromagnetic cortical activity occurring at multiple wavelengths. This data was measured from 100 individuals as part of the Human Connectome Project (Van Essen et al., 2013), and was processed using the Brainstorm software (Baillet et al., 2011) as previously described (Shafiei et al., 2021). Maps were available for the following canonical frequency bands: alpha (8–12 Hz), beta (15-29 Hz), delta (2-4 Hz), theta (5–7 Hz), low gamma (30-59), high gamma (60-90), as well as intrinsic timescale (Gao et al., 2020; Markello et al., 2022). These maps were resampled from Freesurfer 4k space to fslr 32k space using nearest neighbor interpolation, and were parcellated using the Glasser atlas. Regional correlations were conducted between each MEG map and each of the three molecular gradients. The highest correlation for each gradient was visualized and significance was tested using a spatially-aware permutation test. Specifically, an established permutation framework to account for spatial autocorrelation (Seidlitz et al., 2018; Váša et al., 2018) (https://github.com/frantisekvasa/rotate_parcellation) was applied to each of the three molecular gradients to create null brain maps. Each map was correlated to the MEG map with the highest correlation to that gradient, creating a null distribution of r-values. The observed statistic using the true gradient map was compared to this null distribution to derive exact p-values. All correlations between molecular components, MEG maps, and fMRI gradients can be found in **Fig S5**.

### Expression of gradients across development and across species

#### Cross-species gradient expression

We were interested in the degree to which the molecular gradients defined in the Allen would replicate in non-human primate adults, indicating to what degree the gradients are specific to humans. Using the PsychENCODE Human Brain Evolution Adult human, chimpanzee, macaque dataset (see **Replication cohorts**), we calculated correlation values (see **Out-of-sample replication procedure**) for each component across six human, five chimpanzees and five macaque donors, all adults. This resulted in *r*-values for each individual that, for each component, represented the degree to which the individual regional component expression correlated with that of the discovery dataset. We used linear models to evaluate whether there were main effects of species and PLS latent variables, as well as their interaction. To ensure correlations were not a product of the data distribution and low number of tissue samples (n[max]=16), we create individual-specific null effect distribution by, for each individual, permuting their data without replacement and re-running the correlation 100 times. These null distributions were used to define analysis-specific p-values for each individual indicating the likelihood the observed inter-dataset correlation value was achieved by chance given a random set of genes.

We also wanted to ensure that high reproducibility of the gradients was not simply driven by dataset similarity. In other words, we were curious whether the gradient replication is consistent with replication of any coherent biological signal derived from the Allen dataset. To assess this, we created a distribution of biological signals by finding the first 100 principal components in the Allen Human Brain Atlas dataset. Using the same procedure described above (**Replication procedure**), we found correlation values for each individual across all 100 components, and used these values to create individual-specific biological signal distributions. We used these distributions to derive “p-values” for each PLS gradient representing the likelihood that individual-level reproducibility of the gradient exceeds that of the average biological signal (in this case, the average Allen Human Brain Atlas principal component).

#### Developmental gradient expression

We were interested in tracking the emergence of each gradient during brain development, and determining approximately at what developmental epoch these gradients demonstrated adult-like regional expression. In the Brainspan dataset (see **Replication cohorts**), we once again derived correlations for each individual representing similarity to the discovery dataset. In this case, since the discovery dataset was composed of only adult brains, this correlation value can be thought to represent the degree to which gradient expression in the individual resembles average adult gradient expression. These donor-level correlation values were then plotted against log age. To remove 0s, before log transformation, age in both GTEx and Brainspan age was calculated in weeks with week 0 equal to conception (assuming a 40 week pregnancy). To ensure results were interpretable with respect to true age, samples were divided into age groups: fetal (prenatal), infant (0-2 years), child (2-10 years), adolescent (11-19 years) or adult (20+ years). To ensure the observed developmental trajectories were generalizable across datasets, and to evaluate whether they were generalizable across primate species, we repeated the procedure described above for individual human and macaque brains from the PsychENCODE “Developmental rhesus and human dataset” (see **Replication cohorts**). For macaques, weeks start at conception assuming a 24 week gestation period. Age groups for macaques were constructed as follows: fetal (prenatal), infant (0-1 years), child (2-3.5 years), adolescent (3.5-7 years), adult (8+ years).

### Annotation of gradient-associated genes

#### Gene list curation

Genes were identified as significant contributors to each gradient. Significant contribution was defined as the top and bottom 0.83% genes (n=260) of each gradient’s gene list, sorted by PLS X loading. This represents the 5% most contributing genes, divided in half (as both tails were assessed), then divided by three (for multiple comparisons, as three components were assessed). This produced six different gene sets, but for most analyses, top genes were combined across all three components. This gene set is referred to throughout the manuscript as “gradient-associated genes”. Each gene was further categorized based on whether it was part of only one component’s (both tails) gene sets, was present in two components gene sets, or was present in all three.

#### Gene enrichment analysis

Two gene ontology enrichment analyses were conducted. The first evaluated the enrichment of a combination of all 780 top gradient-associated genes. The second analyses involved separate analyses of three different gene sets defined by whether genes contributed to one, two or all three components (see above). For each analysis (four in total), gene lists were submitted as gene sets to ToppGene’s ToppFun enrichment feature (https://toppgene.cchmc.org/enrichment.jsp), using the whole list of Allen Human Brain Atlas genes as the background gene set. Only the following term categories were assessed: GO: Molecular Function, Go: Biological Process, Go: Cellular Component, Pathway (all), Disease (all), and Human Phenotype. All other settings were left to their defaults. Note that ToppGene databases are continuously updated; this ToppGene query was conducted on June 13, 2022. The gene set enrichment analysis further nominated associations with several psychiatric diseases. To validate this finding, we used an established gene-set analysis method (MAGMA) (de Leeuw et al., 2015) to evaluate the enrichment of gradient-associated genes against a published genome-wide association study identifying two distinct factors across multiple psychiatric conditions (Mallard et al., 2022).

#### Post-hoc morphogen analysis

The gene set enrichment analysis revealed several terms relating to brain development. We also had an a priori hypothesis that the gradients should include developmental morphogens (see **Introduction**). Therefore, we identified a recently published list of genes expressed in a gradient-like pattern in the ventricular zone of the developing murine brain (Ypsilanti et al., 2021). We counted the proportion of these genes that overlapped with gradient-associated genes from our study. A p-value for enrichment was calculated by comparing this overlap with that of 1000 permuted gene sets (without replacement) of the same length as the gradient-associated gene set (n=780).

#### mmQTL-based PheWAS

We wished to further explore the relationship between gradient-related genes and aspects of function and behavior. Cross-referencing of gradient-associated genes was performed against a database of multi-ancestry meta-analytic quantitative trait loci (mmQTL), which had systematically been linked to phenotypes via phenome-wide association studies (PheWAS) (Zeng et al., 2022). Associated files were downloaded from Synapse (https://www.synapse.org/#!Synapse:syn23204884/wiki/606411). This data represented SNPs that were i) associated with both brain expression of a gradient-related gene and ii) genome-wide significantly related to a trait. We identified gradient-related genes from this data resource and created a Sankey diagram to summarize two-way associations.

#### Identifying candidate early developmental regulators of spatiomolecular gradients

Despite discovering the spatiomolecular gradients using data from adult human brains, we hypothesize that these signatures reflect changes instantiated during early development. Therefore we used a data-driven approach to identify and validate genes that show regional patterns resembling those of the spatiomolecular gradients already during early brain development (**Fig S7B**). We hypothesized that a proportion of gradient-associated genes would show regional expression patterns resembling the LVs throughout development, including during prenatal periods. Using the Brainspan data as the discovery dataset, we repeated analyses described in Methods section “**Developmental gradient expression”** to derive regional similarity to adult (AHBA) gradients, but for each LV-associated gene across each donor. This process resulted in a developmental trajectory of gradient similarity for each LV-associated gene. For each LV, these data were then clustered using agglomerative clustering with ward criterion (**Fig S7B**). Clustering was repeated for k = 2 to 50, and silhouette index was derived from each clustering solution. Peaks in silhouette index across solutions were identified and examined visually by averaging genes within the same cluster and plotting their expression across age (**Fig S7B**). The lowest k solutions to demonstrate a cluster with high regional similarity to its LV throughout the measurement period (e.g. a non-transitional developmental pattern (Li et al., 2018)) was selected (**Fig S7B**). A k=12 solution was selected for LV1, a k=7 solution for LV2, and a k=17 solution for LV3.

Next, a technique was implemented to “apply” the clustering solution derived in brainspan to the PsychENCODE human data. For selected clustering solutions, cluster centroids were derived for each cluster. This was accomplished by plotting the mean expression of genes within the cluster against log age, and then deriving the fitted lowess curve summarizing this relationship. Next, the fitted lowess curve of expression vs age was derived for each gradient-associated gene in the PsychENCODE dataset. For each gene, mean absolute distance was calculated between its fitted curve and that of each cluster centroid. The gene was then assigned to the cluster for which the shortest distance was observed between the gene’s curve and centroid curve. After this process was repeated for all genes, genes were identified that fell into the non-transitional cluster in both datasets (**Fig S7B**). This set of genes was further curated by eliminating any gene that showed a negative regional correlation with adult LV expression across any samples, and eliminating any gene that never showed a regional correlation with adult LV above 0.5 across any sample. This process resulted in eight genes for LV1, 11 genes for LV2 and 1 gene for LV3 that showed high regional similarity to adult LV expression across brain development.

Finally, we wished to assess whether genes identified in the previous analysis also showed consistent adult LV-like expression throughout development in macaques. This required us to map macaque developmental time to human developmental time, which was accomplished using harmonized developmental periods (Kang et al., 2011; Zhu et al., 2018). We then visualized regional similarity to LV expression across developmental periods for each of the 20 genes identified from the previous analysis across all three datasets (Brainspan, PsychENCODE Macaque, PsychENCODE Human).

## Data and code availability

All analyses in this manuscript were performed using pre-existing datasets, and each of which can be accessed online (see Methods). All figures, tables and analyses can be reproduced using a set of Jupyter notebooks and supplemental scripts: https://github.com/PennLINC/Vogel_PLS_Tx-Space.

## Supplementary Figures

**Fig S1.**
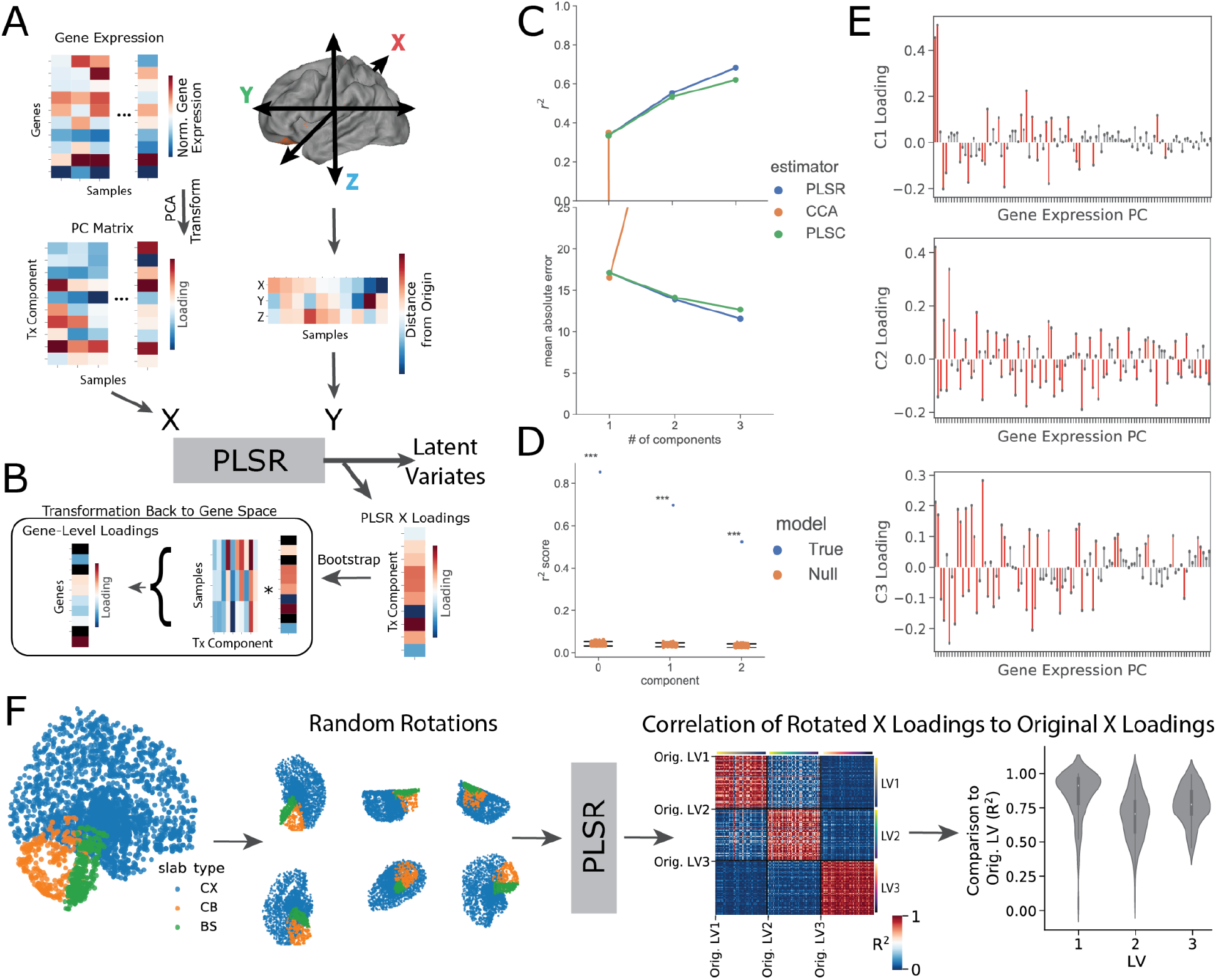
PLS methods schematic. **A)** PLS model was fit to multivariate relationships between gene expression and space. One side of the PLS (PLS X), a gene-by-sample matrix, was decomposed into 100 principal components. The other side (PLS Y) contained x (medial-lateral), y (anterior-posterior) and z (dorsal-ventral) MNI stereotaxic coordinates where x coordinates were converted to absolute values. The resulting PLS components are made up of latent linear combinations of gene expression (PLS X) and spatial positioning (PLS Y) that demonstrate maximal covariance (as in **Fig 1**). **B)** The dot product of the PLS rotation vector and the transposed standardized principal component matrix from A) was found, resulting in PLS X loadings in gene space. **C)** In the training set, 100 rounds of 10-fold cross-validation were used to optimize performance over three different estimators (PLS-R, PLS-C, CCA) and over number of PLS components (matrix rank for PLS Y in this case is 3, so C=1-3 evaluated). Point plots show R^2^ (top) and MSE (bottom) across estimators (hue) and number of components (x-axis). Confidence intervals are present but not visible in the plot. CCA performed poorly when number of components > 1, whereas PLSC and PLSR showed improved performance with increasing components. Top performance was achieved using PLSR with 3 components. **D)** The optimal model from C was rerun 1000 times with shuffled labels and explained variance of each component was recorded. This null distribution (orange dots) was used to determine if the explained variance of the empirical PLS model (blue dot) was greater than what would be expected by chance given the dataset. **E)** PLS X loadings for all 100 input features (gene expression principal components) for each of the 3 PLS latent variables summarized over 1000 bootstrap samples. Red-colored features indicate that the 95% CI of the mean feature weight across bootstrap samples did not cross 0, indicating good reliability. Importantly, PC1 (the first feature) did not dominate any PLS LV and was distributed fairly evenly across all 3 PLS LVs, indicating none of the PLS LVs are simply equivalent to PC1. **F)** To ensure molecular gradients were not driven by arbitrary spatial coordinates, (left) the brain was rotated 100 times and, for each rotation, the same gene expression by (rotated) x,y,z coordinate PLS was run. (right) The loadings of each set of X (gene expression) LVs were correlated. The correlation matrix shows the correlation (R2) between the X loadings of all three LVs for each rotation, as well as for the original (orig.; i.e. non-rotated) PLS model. The matrix is ordered first by LV, then by rotation, and clearly shows high within-LV and low between-LV correlations across rotations. The violin plot shows that the X loadings across rotations were highly consistent with those of the original PLS model. This indicates that spatiomolecular gene-expression gradients were not driven by base brain orientation.

**Fig S2.**
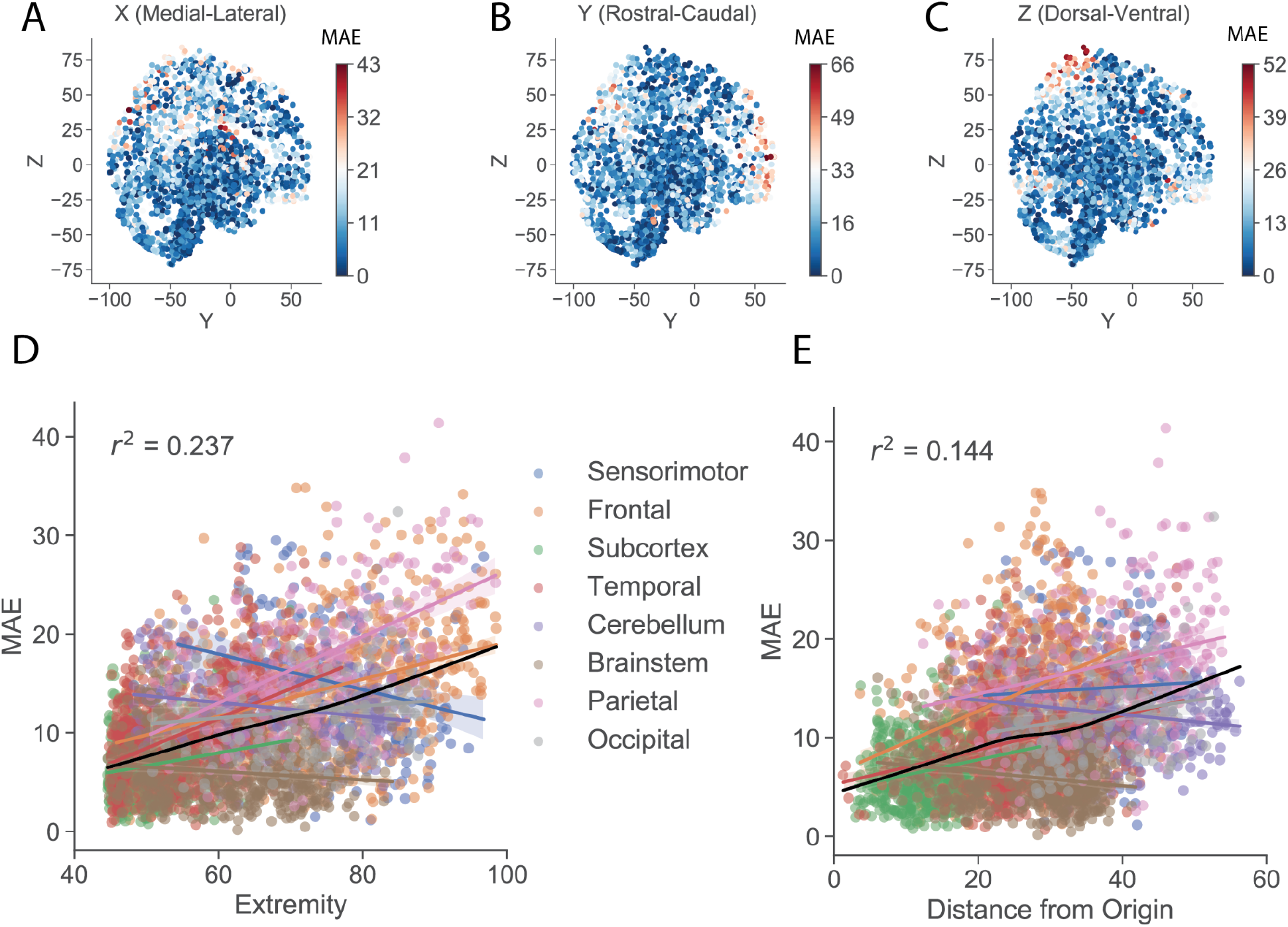
Quantifying error in PLS model. Figure 1 shows the overall absolute error associated with each sample. Here, error specifically within **A)** x-, **B)** y- and **C)** z-coordinates are visualized. **D)** Extremity of each sample was quantified as the average distance from all other samples. Extremity explained approximately 24% of the variance in sample error, such that the location of samples extracted from more extreme spatial locations of the brain were less accurately predicted. This relationship was driven especially by samples located in frontal, parietal and temporal neocortex. **E)** Similar to D), the location of samples located further from the origin (MNI coordinate 0,0,0) were more poorly predicted by the PLS model.

**Fig S3:**
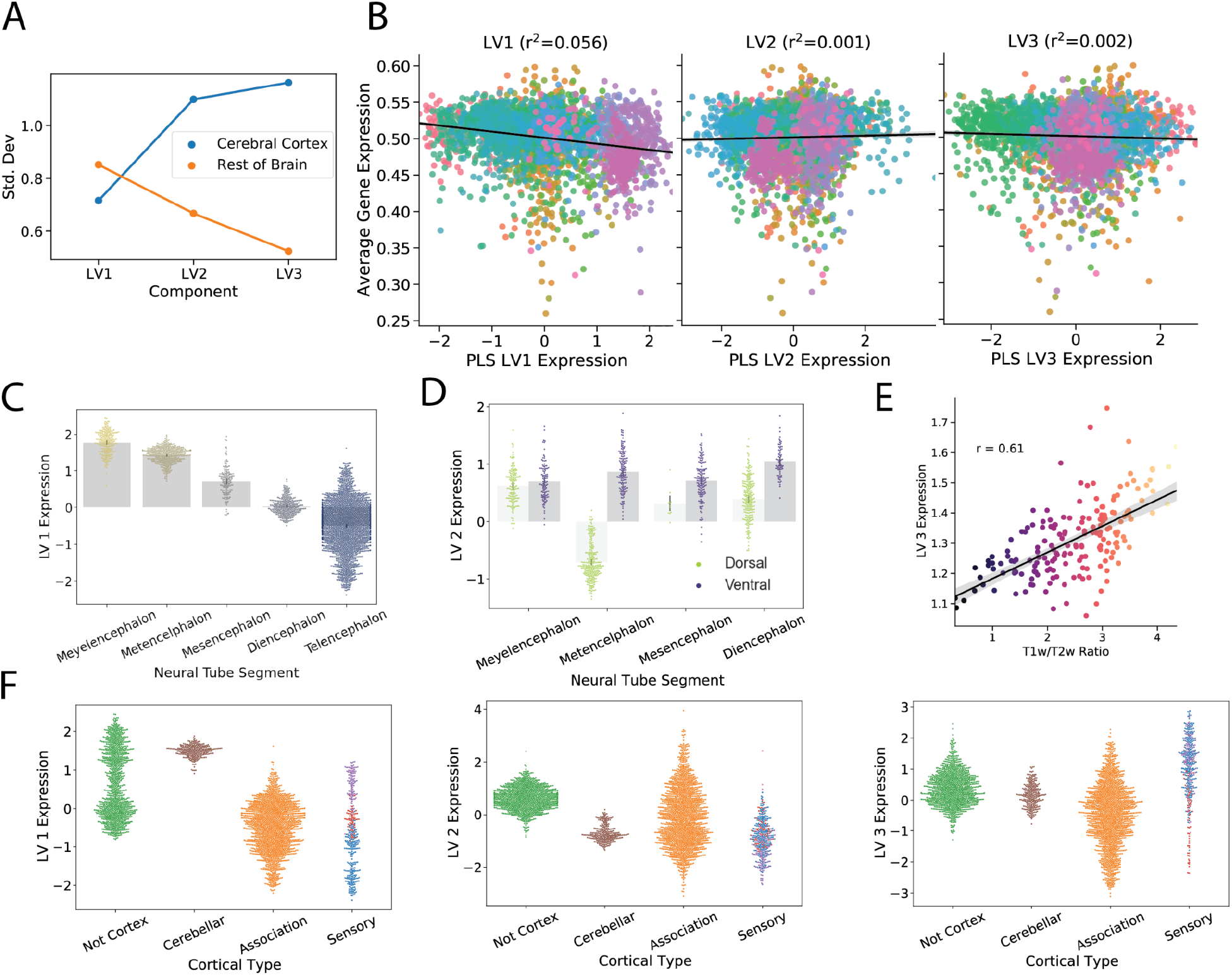
Spatiomolecular gradients vary systematically with fundamental developmental and cortical organizational features. **A)** Total signal variation of each component across the cerebral cortex and non-cortex, represented as the standard deviation of each component across regions. LV3 showed the greatest variation in the cerebral cortex and the least outside of it. **B)** Expression of each component correlated against total gene expression averaged across genes. Each dot represents a brain region. Dots are colored in accordance with brain division. Plots demonstrate that PLS latent variables are not simply driven by overall regional variation in gene expression. Note also that regional variation varies strongly across the X but not Y axis of each plot. **C)** Regions were divided into categories based on from which developmental compartment they originated. LV1 expression demonstrated a linear rostral-caudal gradient along developmental compartments. **D)** Within each compartment, regions were further divided based on whether they originated from dorsal or ventral aspects of the developing human brain (this information is not available for the telencephalon). Across compartments, LV2 expression differed between dorsal- and ventral-originating regions. **E)** Cortical LV3 expression correlates (r=0.78) with the T 1w/T2w ratio, an MRI-derived index of myelination. **F)** All regions were categorized based on whether they were part of sensory cortex, association cortex, cerebellar cortex or non-cortex (i.e., everything else). The distribution in expression of LV1 (left), LV2 (middle) and LV3 (right) across these cortical types is visualized. Each component carries a distinct cortical profile: LV1 expression is low in cerebral cortex but high in other types, LV2 expression is low in all cortical types but high in non-cortex, and LV3 expression is low in association cortex but high in all other cortical types. These results suggest that cortical organization is organized along the three molecular axes.

**Fig S4:**
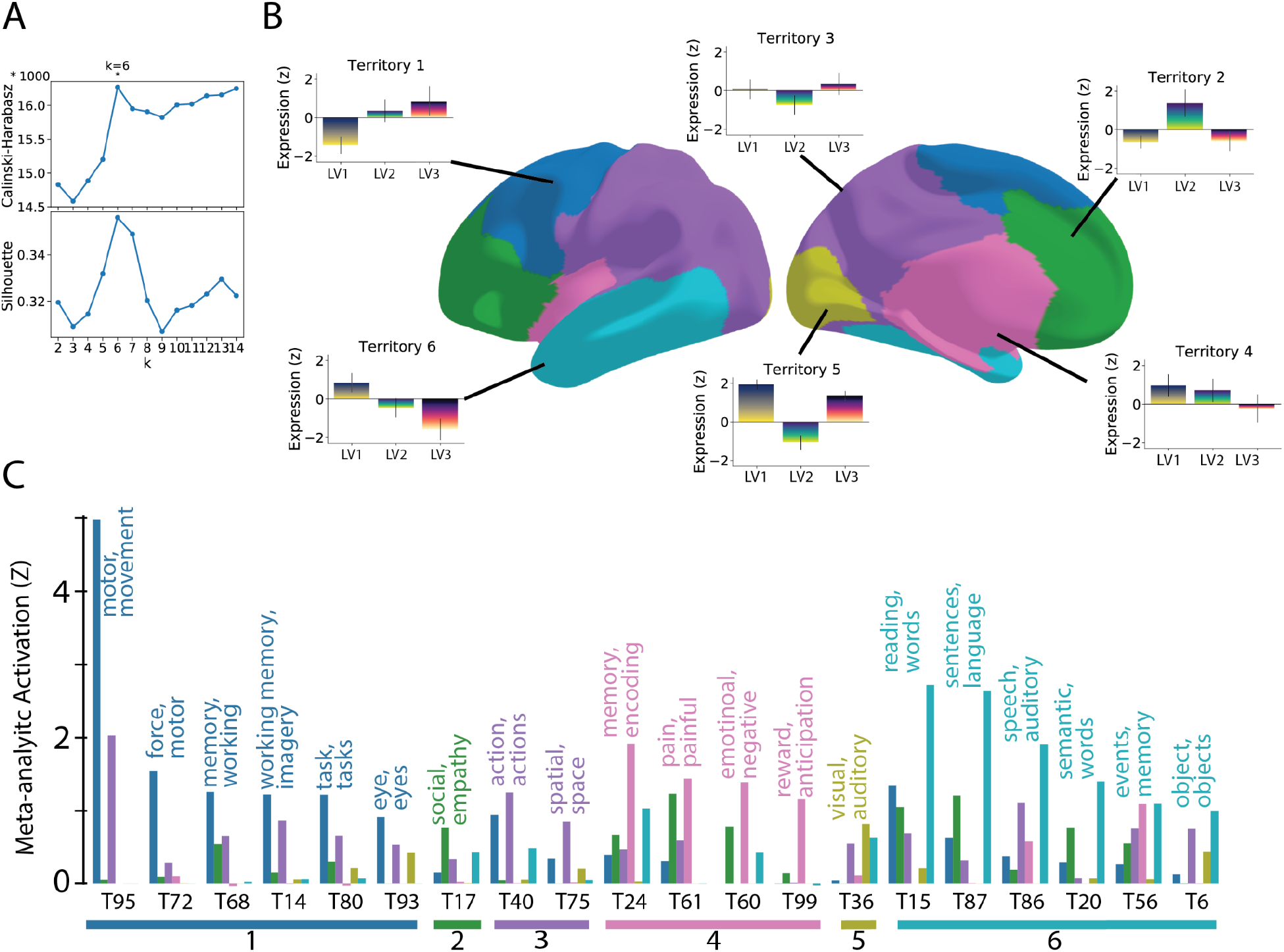
The interaction of functional gradients in the cerebral cortex creates functionally distinct territories. **A)** Vertex-wide clustering of PLS LV concentrations suggested six molecular territories, based on consensus from common clustering metrics. **B)** Vertex labels of each cluster representing six molecular territories composed of different concentrations of PLS gradients. Surrounding bar plots indicate relative concentrations of each gradient within each territory. **C)** Molecular territories form functionally distinct domains: 68 cognition-associated meta-analytic topic maps were downloaded from neurosynth, representing common activation patterns across studies in association with inter-related terms. Average z-scores for each map were computed for each functional territory. Plot shows each term that fell into the top 5 highest-rank z values for one of the six molecular territories, indicating the mean z for each territory, and the top two words associated with the topic map. The x-axis shows the Topic labels, for which definitions are available at neurosynth.org and in **Table S8.**

**Fig S5.**
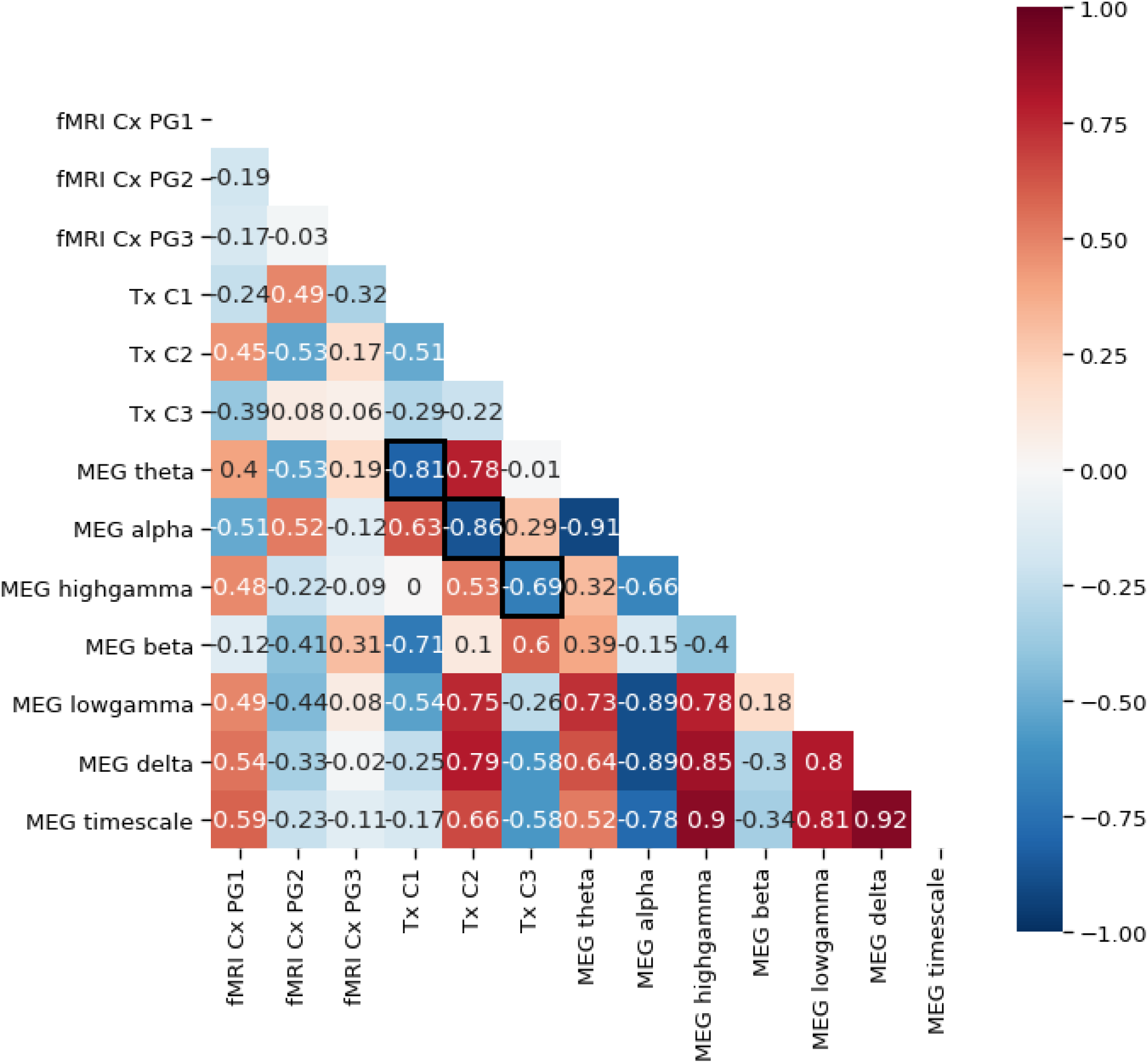
Spatial relationships between molecular and functional gradients of the cerebral cortex. Correlation matrix representing spatial relationship between fMRI principal gradients 1-3 (fMRI Cx PG1-3), the three molecular gradients described in this study (Tx LV1-3), and seven MEG maps (meg alpha, beta, delta, theta, low gamma, high gamma, intrinsic timescale). For each molecular gradient, the strongest relationship is highlighted with bolded box borders. These relationships are visualized in main text **Fig 2F**.

**Fig S6.**
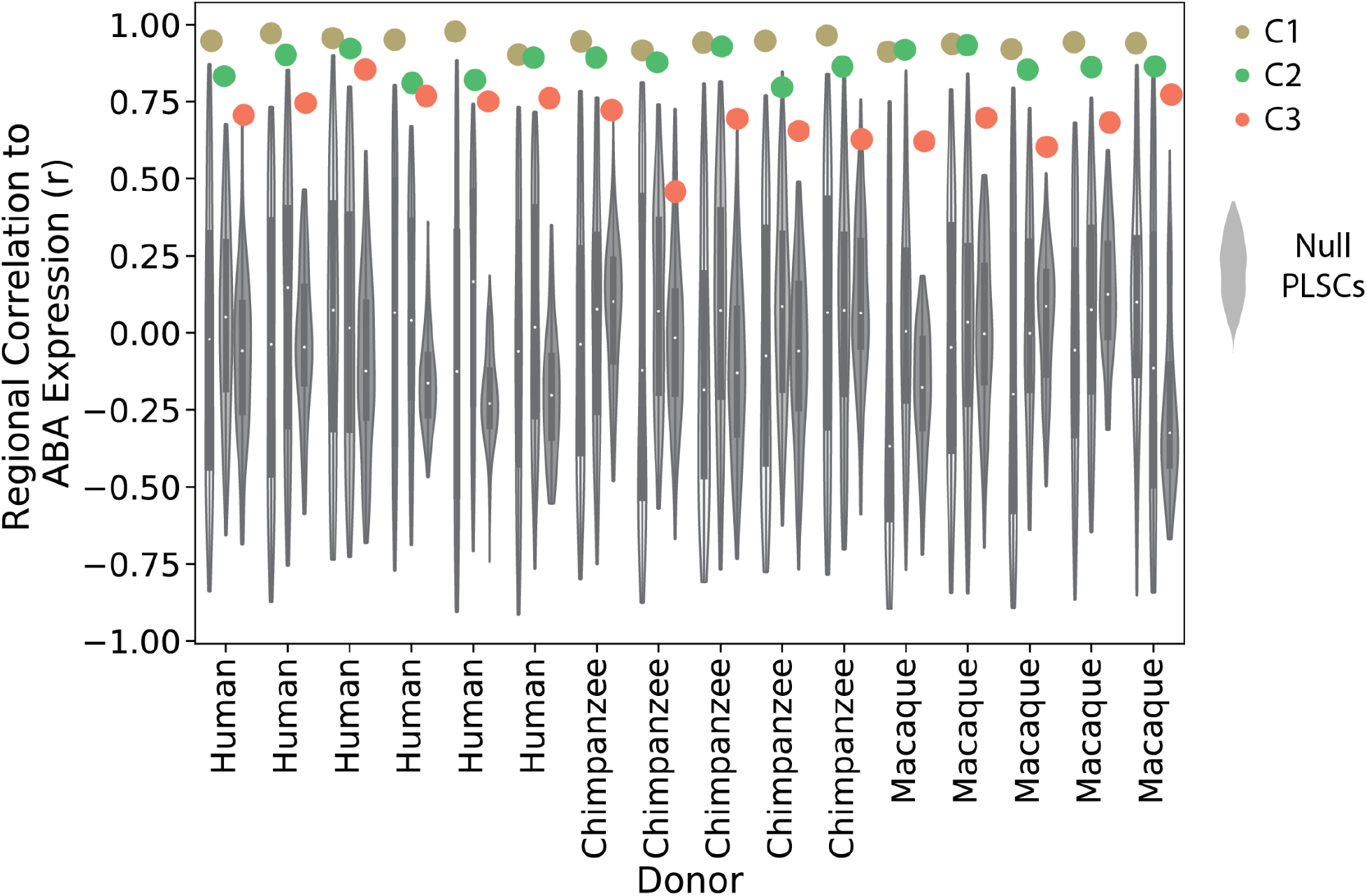
Individual null models for PLS latent variable replication. As described in Figure 4A, six adult human, five adult chimpanzee and five adult macaque brains were available from (http://evolution.psychencode.org; (Zhu et al., 2018)) with tissue samples extracted from the same 16 brain regions. Component correlations (as described in **Fig 1I**) for each LV were generated for each individual. Null models were generated for each individual. Specifically, PLS weights were shuffled 100 times for each component, and the reproducibility analysis (Fig **3A**) was conducted using each of the shuffled weights, separately for each individual. The resulting similarity (r) values were used to create a null distribution specific to each component for each individual. Null distributions are visualized, while the “true” similarity (r) values for LV1 (yellow), LV2 (green) and LV3 (orange) are represented as colored dots. These suggest that the successful replication of the PLS LVs is greater than would be expected given the structure of the data and overall similarity between datasets.

**Fig S7.**
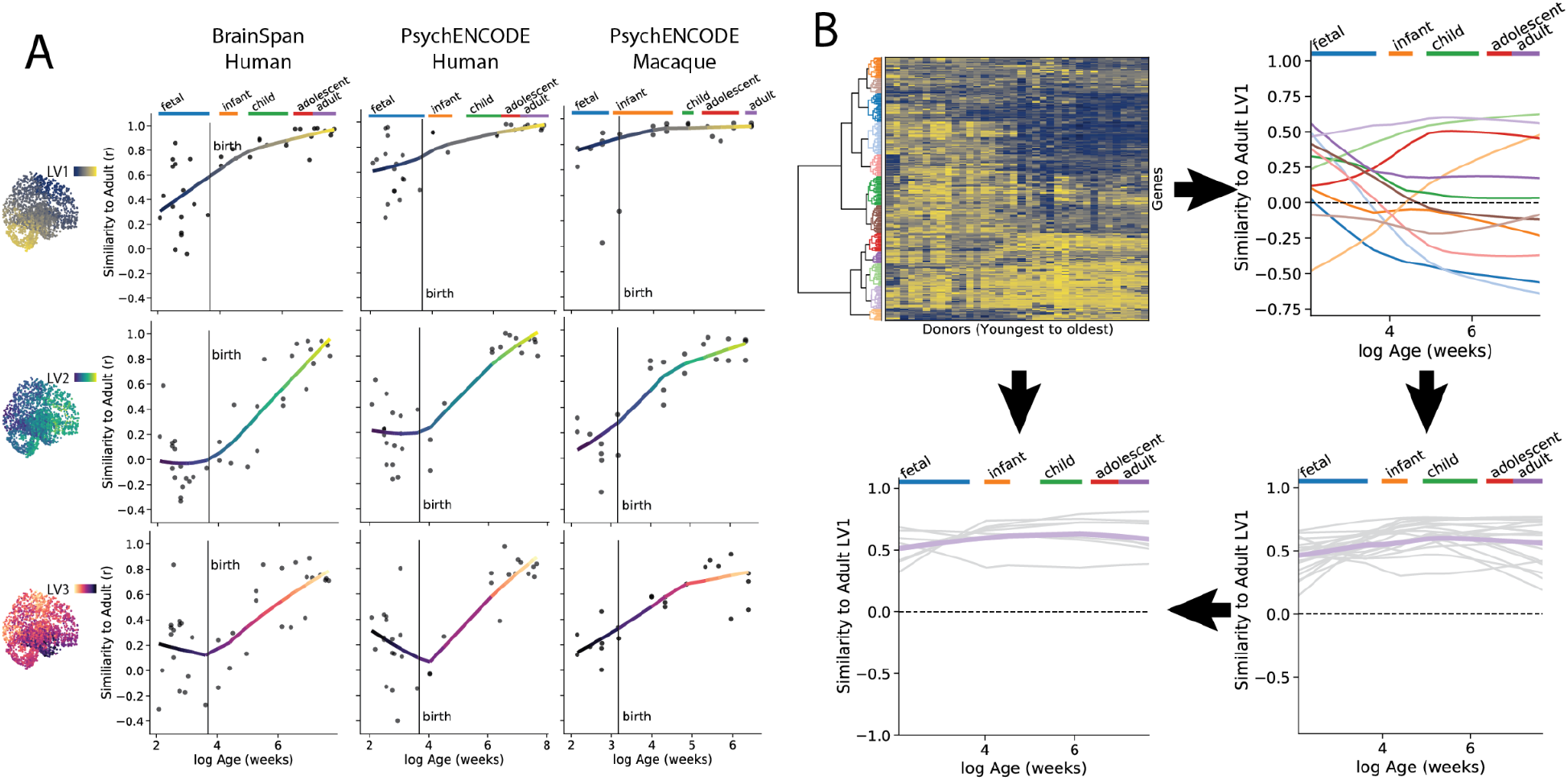
Individual trajectories of PLS latent variables over development. **A)** The same data plotted in Main Text **Fig 3C**, but separated by PLS latent variable. Here, it is possible to appreciate increased variation in component expression in early prenatal stages of human datasets, perhaps suggesting an early-prenatal peak followed by a perinatal dip. **B)** Workflow for gene-trajectory clustering, using LV1 as an example. (Top right) For each brain donor in Brainspan, the regional similarity to LV1 (**Fig 1I**) was calculated for each of the top LV1 genes. This data was then clustered to identify genes with similar developmental trajectories. (Top right) The developmental trajectory of each cluster is visualized, where the x-axis is developmental time (log of age in post-conception weeks) and the y-axis is regional expression similarity to LV1. (Bottom right) One cluster demonstrated a non-transitional trajectory with consistently high regional similarity to LV1. The individual trajectories of each contributing gene is visualized. (Bottom left) The clustering solution from Brainspan was applied to PsychENCODE data to identify genes demonstrating the same trajectory. Genes that fell into the non-transitional cluster across both datasets were considered to be reliable and are highlighted in the main text and **Fig. 3D**.

**Fig S8.**
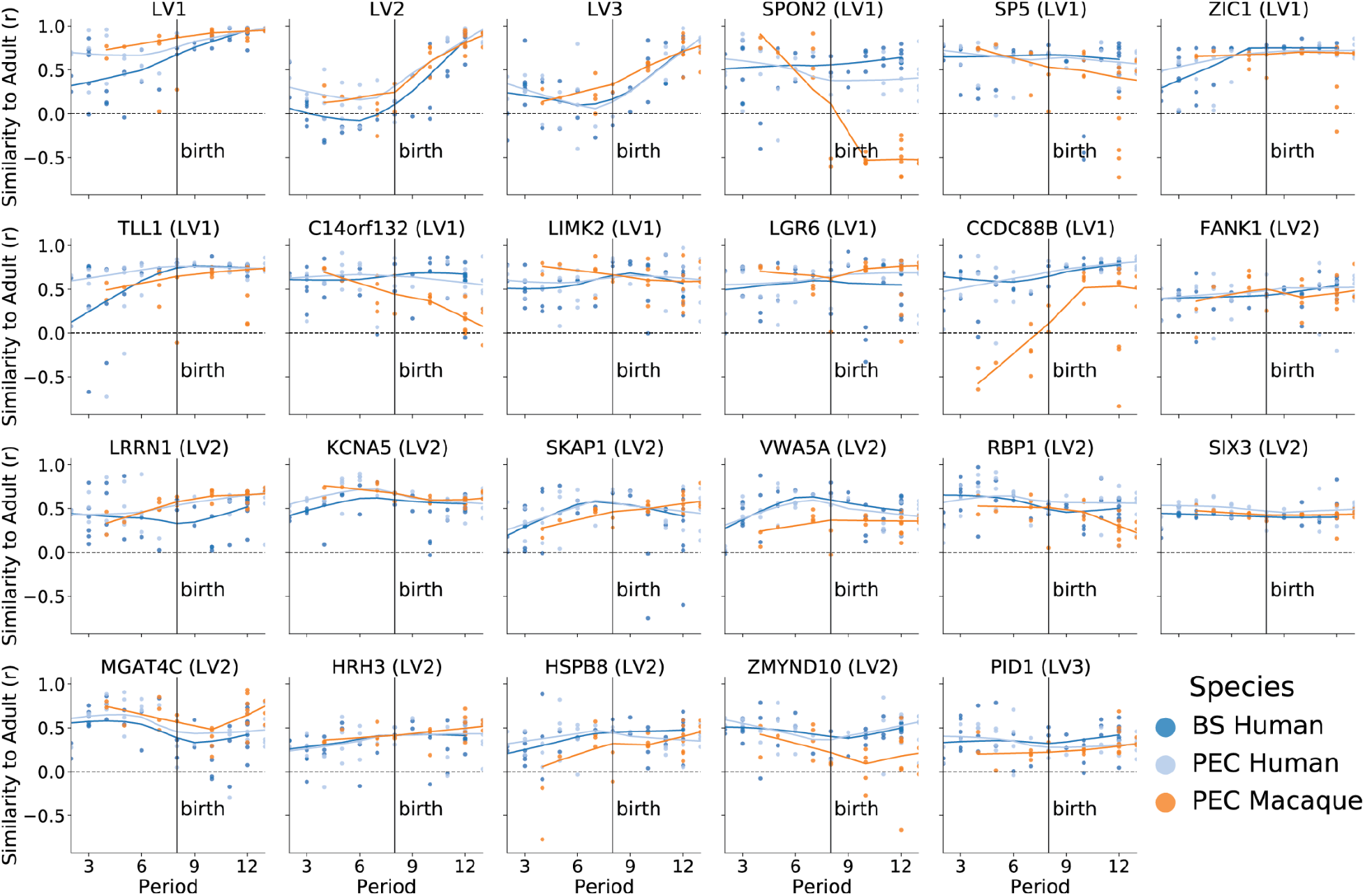
Comparison of candidate LV-maintenance genes across primate species. In order to establish whether genes identified as candidates for LV maintenance were consistent across datasets, it was necessary to align macaque and human developmental timelines to a shared temporal space. We used the developmental periods described in (Kang et al., 2011), and assigned the primate PsychENCODE data to these periods based on (Zhu et al., 2018). For all three LVs and for each of the 21 candidate genes, we visualize regional similarity to adult (AHBA) latent variable expression across brain development. The x-axis of each plot represents developmental period and the y-axis represents similarity to adult regional LV expression. The parenthetical LV in each plot title indicates which LV is being compared. Most genes showed highly similar non-transitional developmental trajectories across species, though there were some notable exceptions (e.g. *SPON2, CCDC88B*).

